# Universal open MHC-I molecules for rapid peptide loading and enhanced complex stability across HLA allotypes

**DOI:** 10.1101/2023.03.18.533266

**Authors:** Yi Sun, Michael C. Young, Claire H. Woodward, Julia N. Danon, Hau Truong, Sagar Gupta, Trenton J. Winters, George Burslem, Nikolaos G. Sgourakis

## Abstract

The polymorphic nature and intrinsic instability of class I major histocompatibility complex (MHC-I) and MHC-like molecules loaded with suboptimal peptides, metabolites, or glycolipids presents a fundamental challenge for identifying disease-relevant antigens and antigen-specific T cell receptors (TCRs), hindering the development of autologous therapeutics. Here, we leverage the positive allosteric coupling between the peptide and light chain (β_2_ microglobulin, β_2_m) subunits for binding to the MHC-I heavy chain (HC) through an engineered disulfide bond bridging conserved epitopes across the HC/β_2_m interface, to generate conformationally stable, open MHC-I molecules. Biophysical characterization shows that open MHC-I molecules are properly folded protein complexes of enhanced thermal stability compared to the wild type, when loaded with low- to intermediate-affinity peptides. Using solution NMR, we characterize the effects of the disulfide bond on the conformation and dynamics of the MHC-I structure, ranging from local changes in β_2_m interacting sites of the peptide binding groove to long-range effects on the α_2-1_ helix and α_3_ domain. The interchain disulfide bond stabilizes empty MHC-I molecules in a peptide-receptive, open conformation to promote peptide exchange across multiple human leucocyte antigen (HLA) allotypes, covering representatives from five HLA-A, six HLA-B supertypes, and oligomorphic HLA-Ib molecules. Our structural design, combined with conditional β-peptide ligands, provides a universal platform for generating ready-to-load MHC-I systems of enhanced stability, enabling a range of approaches to screen antigenic epitope libraries and probe polyclonal TCR repertoires in the context of highly polymorphic HLA-I allotypes, as well as oligomorphic nonclassical molecules.

**Significance Statement:** We outline a structure-guided approach for generating conformationally stable, open MHC-I molecules with enhanced ligand exchange kinetics spanning five HLA-A, all HLA-B supertypes, and oligomorphic HLA-Ib allotypes. We present direct evidence of positive allosteric cooperativity between peptide binding and β_2_m association with the heavy chain by solution NMR and HDX-MS spectroscopy. We demonstrate that covalently linked β_2_m serves as a conformational chaperone to stabilize empty MHC-I molecules in a peptide-receptive state, by inducing an open conformation and preventing intrinsically unstable heterodimers from irreversible aggregation. Our study provides structural and biophysical insights into the conformational properties of MHC-I ternary complexes, which can be further applied to improve the design of ultra-stable, universal ligand exchange systems in a pan-HLA allelic setting.

## Introduction

The proteins of the class I major histocompatibility complex (MHC-I) are essential components of adaptive immunity in all jawed vertebrates(1). They function by displaying a broad spectrum of self, aberrant, or foreign epitopic peptides, derived from the endogenous processing of cellular proteins on the cell surface, thereby enabling immune surveillance by cytotoxic T lymphocytes (CTL) and Natural Killer (NK) cells(2). Classical MHC-I molecules comprise a 8-to-15-amino-acid peptide, an invariable light chain human β_2_ microglobulin (β_2_m), and a highly polymorphic heavy chain (HC) that contains three extracellular domains (α_1_, α_2_, and α_3_)(3). The expansion of MHC-I genes in humans (human leukocyte antigens, or HLAs) has resulted in more than 35,000 alleles with polymorphic residues located on the peptide binding groove, composed by the α_1_ and α_2_ domains and a β-sheet floor(4). The polymorphic nature of HLA allotypes leads to a diversity of displayed peptide repertoires and interactions with molecular chaperones, other components of the antigen processing pathway, and T cell receptors (TCR), which ultimately define immune responses and disease susceptibility. Classical HLA-A and HLA-B allotypes can be further classified into 12 supertypes according to their various peptide binding specificity, determined by primary anchoring interactions with the peptide positions 2, 3, 5, and 9(5, 6). Therefore, recombinant peptide-loaded MHC-I (pMHC-I) molecules are typically generated using *in vitro* refolding together with synthetic peptides(7) as soluble monomers and can be further prepared as tetramers or multimers(8). These reagents have been one of the most important tools for detecting, isolating, and stimulating CTLs, and screening, optimizing, and identifying immunodominant T cell epitopes for immunotherapy, diagnosis, and vaccine development(9).

The assembly and peptide loading of nascent pMHC-I molecules occurs in the lumen of the endoplasmic reticulum and involves many molecular chaperones, including the peptide loading complex (PLC)-restricted tapasin and the PLC-independent homologous, transporter associated with antigen processing (TAP)-binding protein related (TAPBPR). The folding of the MHC-I HC and the formation of disulfide bonds in the α_2_ and α_3_ domains is assisted by calnexin and ERp57(10). The HC then assembles with β_2_m to generate an empty heterodimer that is highly unstable for most MHC-I alleles, which is stabilized by association with tapasin, ERp57, calreticulin, and TAP in the PLC(11). Both chaperones tapasin and TAPBPR can facilitate the binding of high-affinity peptides to confer the stability and proper trafficking of MHC-I, which finally resides on the cell surface for hours to days(12). Loading of high-affinity peptides induces a “closed” conformation of the α_2-1_ helix via negative allosteric modulation between non-overlapping peptide binding sites and tapasin/TAPBPR binding sites to release the chaperones(13). However, peptide loading and β_2_m binding to the HC are positively allosterically coupled, together stabilizing the ternary complex(14). The peptide loading process is initiated by a rate-limiting step of β_2_m association with the HC, which yields HCs with enhanced peptide binding affinity in the nanomolar range(15, 16). Consequently, maintaining a stable tertiary structure of pMHC-I requires not only the selection of a high-affinity peptide but also the proper function of β_2_m as a conformational chaperone(14, 15). Although under sub-physiological temperatures, stable, peptide-deficient MHC-I HC/β_2_m heterodimers have been reported to express on the cell surface, empty MHC-I molecules have a short half-life, and are rapidly internalized(17, 18). This intrinsic instability of empty MHC-I causes non-specific exogenous peptide binding and irreversible denaturation *in vitro*, limiting its application as an off-the-shelf molecular probe for ligand screening and T cell detection.

A tremendous amount of work has been invested in the biophysical characterization and engineering of pMHC-I molecules, with specific efforts being made to understand the molecular mechanisms of peptide loading, and to develop tools for peptide exchange. Conditional ligands, bound to the MHC-I, can be cleaved by UV exposure or released by increasing the temperature to generate empty molecules, which can be loaded with a rescuing peptide(19–21). Dedicated MHC-I chaperones, including TAPBPR and its orthologs, have also been used to stabilize an array of different MHC-I allotypes in a peptide-receptive conformation with a preferred binding to HLA-A over HLA-B and HLA-C alleles, promoting the exchange of low- to intermediate-affinity peptides for high-affinity peptides in a process known as “peptide-editing”(22–26). In addition, molecular dynamics (MD) simulations combined with a tryptophan fluorescence assay have shown that empty MHC-I molecules are not molten globules like previously reported(27), but have varying degrees of structure in the α_1_ and α_2_ helices(28, 29). More recent studies aim to stabilize peptide-deficient molecules by introducing a disulfide bond across the α_1_ and α_2_ helices, restricting the highly flexible F pocket of the peptide binding groove to mimic the peptide-bound state for common alleles, such as HLA-A*02:01, HLA-A*24:02, and HLA-B*27:05(30–34). Other studies have sought to stabilize the pMHC-I complex by characterizing the interaction of mutant or orthologous β_2_m variants. A functional study using human and murine β_2_m variants bound to HLA-A*02:01 and H-2D^b^ demonstrated that human β_2_m has a greater affinity for H-2D^b^ than murine β_2_m, resulting in enhanced complex stability due to a marked increase in polarity and the number of inter-chain hydrogen bonds(35). Another study identified a S55V mutant β_2_m characterized by its capability to stabilize pMHC-I molecules to a greater extent than the wild type (WT), enhancing peptide binding and CD8+ T cell recognition(36). These studies, altogether, have emphasized the importance and the possibility of generating conformationally stable, peptide-receptive MHC-I heterodimers across different allotypes by manipulating the malleable HC/β_2_m interface.

In this work, we outline an alternative, structure-guided approach to engineering conformationally stable, peptide-receptive, open MHC-I molecules by introducing a disulfide bond bridging conserved sites across the HC/β_2_m interface. We exploit the allosteric mechanisms governing the assembly of MHC-I complexes by locking pMHC-I proteins in an open, peptide-receptive state via the introduction of G120C and H31C mutations on flexible loop regions of the HC and β_2_m, respectively. We show that the interchain disulfide bond increases the thermostability of molecules loaded with low- and moderate-affinity peptide cargo. We use solution nuclear magnetic resonance (NMR) and hydrogen-deuterium exchange mass spectrometry (HDX-MS) to characterize a peptide-receptive, open conformation, and further demonstrate that engineered open MHC-I molecules have improved peptide exchange efficiency and overall stability across five HLA-A, all HLA-B supertypes(37), oligomorphic HLA-Ib alleles, HLA-E, -F and -G, and the MHC-like molecule MR1. Finally, we demonstrate that open MHC-I molecules are functionally competent in detecting antigen-specific cell populations, serving as a universal platform for identifying immunodominant peptide epitopes and probing T cell responses in research, preclinical or diagnostic settings.

## Results

### Structure-guided disulfide engineering stabilizes suboptimal MHC-I ligands

To engineer stable HLA molecules across different allotypes for rapid peptide exchange, we aimed to bridge the HC and the light chain β_2_m through a disulfide bond based on the positive cooperativity between peptide and β_2_m association with the HC(35, 38). We utilized a structure-guided approach by first aligning 215 high-resolution pMHC-I crystal structures that were curated in our previously developed database, HLA3DB (**Fig. 1A**). We found an average distance of 4.25 Å (3.7 Å ≤ Cβ-Cβ ≤ 4.9 Å) between positions G120 of the HC and H31 of the β_2_m (**Fig. 1A**). The distances between the paired residues G120 and H31 on the HC and the β_2_m, respectively, fall within the molecular constraints (5.5 Å) for disulfide cross-linkage(39, 40). The structure of HLA-A*02:01/β_2_m shows that both regions are composed of flexible loops (**Fig. 1B**), which increase the probability of the two cysteine mutations forming a 90° dihedral angle necessary for disulfide bond formation(41). Additionally, a sequence alignment using 75 distinct HLA allotypes with a greater than 1% global population frequency revealed a conserved glycine at position 120, suggesting a potential generality of the design across distinct HLA allotypes, covering various HLA supertypes that can present diverse peptide repertoires (**Fig. S1**). Selected residues G120 and H31 between the HC and β_2_m were further computationally validated using Disulfide by Design(42). Together, the structure and sequence alignments indicate the possibility of applying the interchain disulfide cross-linkage to a broad range of HLA allotypes, including oligomorphic HLA-Ib and monomorphic nonclassical MHC-I-related proteins, to stabilize their ligand-receptive conformations.

**Figure 1.**
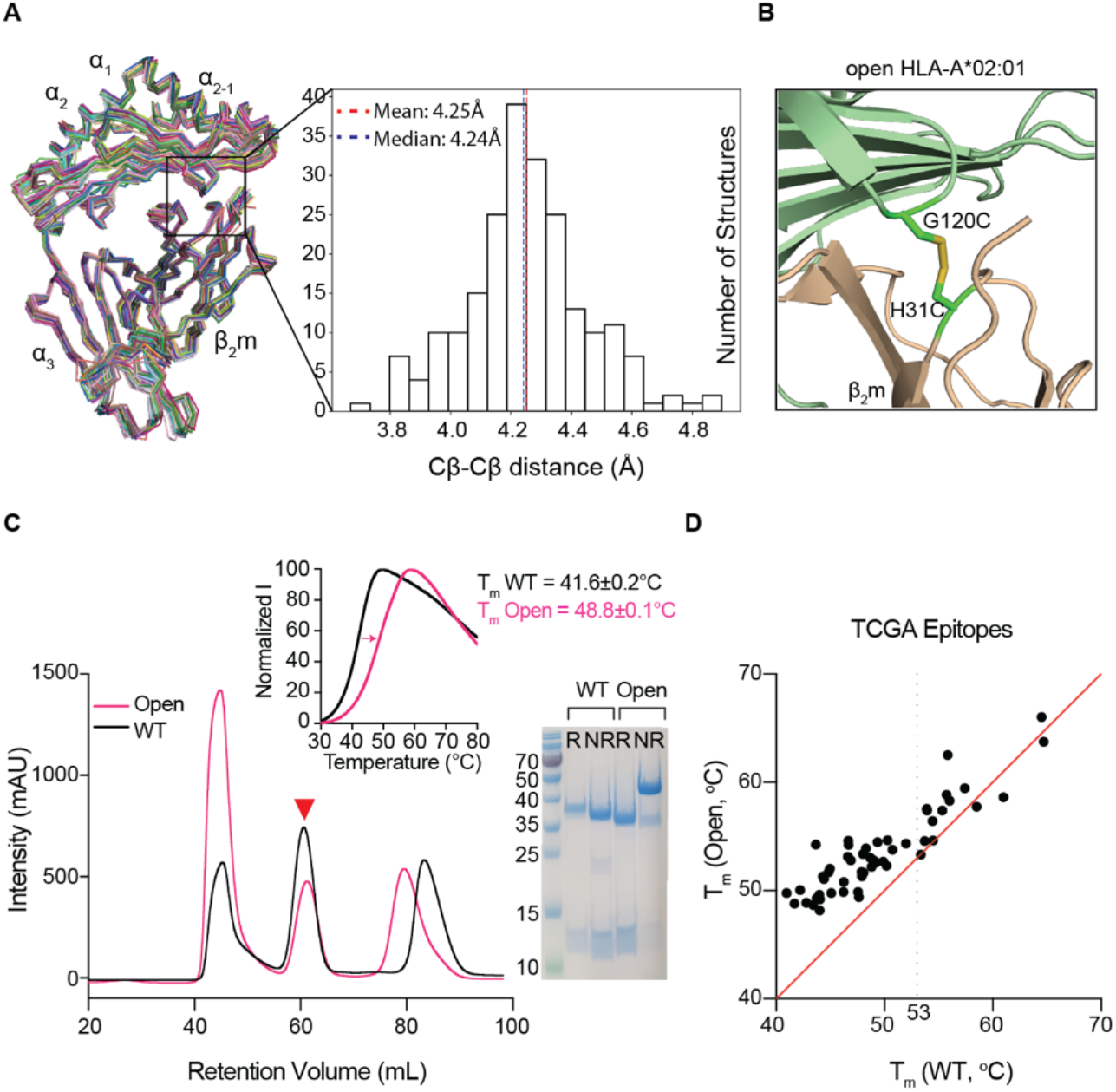
Structure-guided stabilization of suboptimal peptide-loaded HLA-A*02:01 by engineered disulfide between the HC and β_2_m. **A.** Structure alignment and distribution of Cβ-Cβ distances between positions G120 of the HC and H31 of the β_2_m derived from 215 pMHC-I/β_2_m co-crystal structures with resolution values ≤ 3Å. The structures of 52 distinct alleles are aligned by Cα atoms of α_1_, α_2_, and α_3_ domains as ribbons. **B.** Structural model of HLA-A*02:01/β_2_m/TAX9 (PDB ID:1DUZ) with G120 and H31 mutated to cysteines. HLA-A*02:01 HC was colored in light green and β_2_m in wheat. **C.** SEC traces of the WT (black) and the G120C/H31C open (pink) HLA-A*02:01/β_2_m/TAX8. The red triangle arrowhead indicates the complex peaks and is further confirmed by SDS/PAGE analysis in reduced (R) or non-reduced (NR) conditions. DSF shows thermal stability curves of the WT in black (T_m_ = 41.6°C) and the open variant in pink (T_m_ = 48.8°C). The average of three technical replicates (mean) is plotted. **D.** Thermal stabilities correlation of the WT and open HLA-A*02:01/TAX8 loaded with each of 50 peptides from the Cancer Genome Atlas (TCGA) epitope library are shown in dots. The average of three technical replicates (mean) is plotted. The red line represents a conceptual 1:1 correlation (no difference in thermal stabilities).

We next sought to validate the design experimentally by expressing the G120C variant of one of the most common alleles HLA-A*02:01 in *Escherichia coli*, isolated denatured proteins from inclusion bodies, and refolded it *in vitro* with the H31C variant of the β_2_m in the presence of a low-affinity placeholder peptide, TAX8 (LFGYPVYV). Size exclusion chromatography (SEC) and SDS/PAGE confirmed the formation of a G120C/H31C HLA-A*02:01/β_2_m complex (hereafter referred to as open HLA-A*02:01) and the interchain disulfide bond (**Fig. 1C**). We then performed differential scanning fluorimetry (DSF) and observed a substantial improvement in the thermal stability of the open HLA-A*02:01 compared to the WT with melting temperatures (T_m_) of 48.8 °C and 41.6 °C, respectively (**Fig. 1C**). Furthermore, the WT and open HLA-A*02:01/β_2_m/TAX8 were further exchanged with 50 peptides from the Cancer Genome Atlas (TCGA) epitope library (**Table S1**). While the resulting T_m_ values of WT and open HLA-A*02:01 loaded with high-affinity peptides (WT T_m_ ≥53°C) were similar, a pronounced stabilizing effect was demonstrated on the open over WT HLA-A*02:01 when suboptimally loaded with low-to moderate-affinity peptides (WT T_m_ < 53 °C) (**Fig. 1D**). Therefore, evaluation of thermal stabilities for HLA-A*02:01-restricted epitopes spanning a broad range of affinities showed that disulfide linkage between the HC and β_2_m did not impede peptide binding, and consistently support the role of β_2_m in chaperoning and stabilizing the HC for peptide loading.

### Disulfide-engineered pMHC-I shows conformational changes at dynamic sites

Conformational plasticity and dynamics have been previously shown to be important for several aspects of MHC-I function, including peptide loading, chaperone recognition, and TCR triggering(43–46). To elucidate differences in conformational landscapes of peptide-loaded MHC-I molecules, we used established solution NMR methods(47). First, we refolded WT and open A*02:01/β_2_m complexes with a high-affinity MART-1 peptide ELAGIGILTV, which were isotopically labeled with ^15^N, ^13^C, and ^2^H at either the MHC-I heavy or the β_2_m light chain, followed by reintroduction of exchangeable protons during complex refolding. After independently assigning both protein subunits using a suite of TROSY-based triple resonance experiments(48) (**Fig. S2, S3**), we measured differences in backbone amide chemical shifts between the WT and open HLA-A*02:01. We then calculated chemical shift perturbations (CSPs) capturing both the amide ^1^H and ^15^N chemical shift changes. Residues showing CSPs above 0.05 ppm (5 times the sensitivity relative to ^1^H) were mapped on the complex structure to highlight sites undergoing changes in the local magnetic environment (**Fig. 2**).

**Figure 2.**
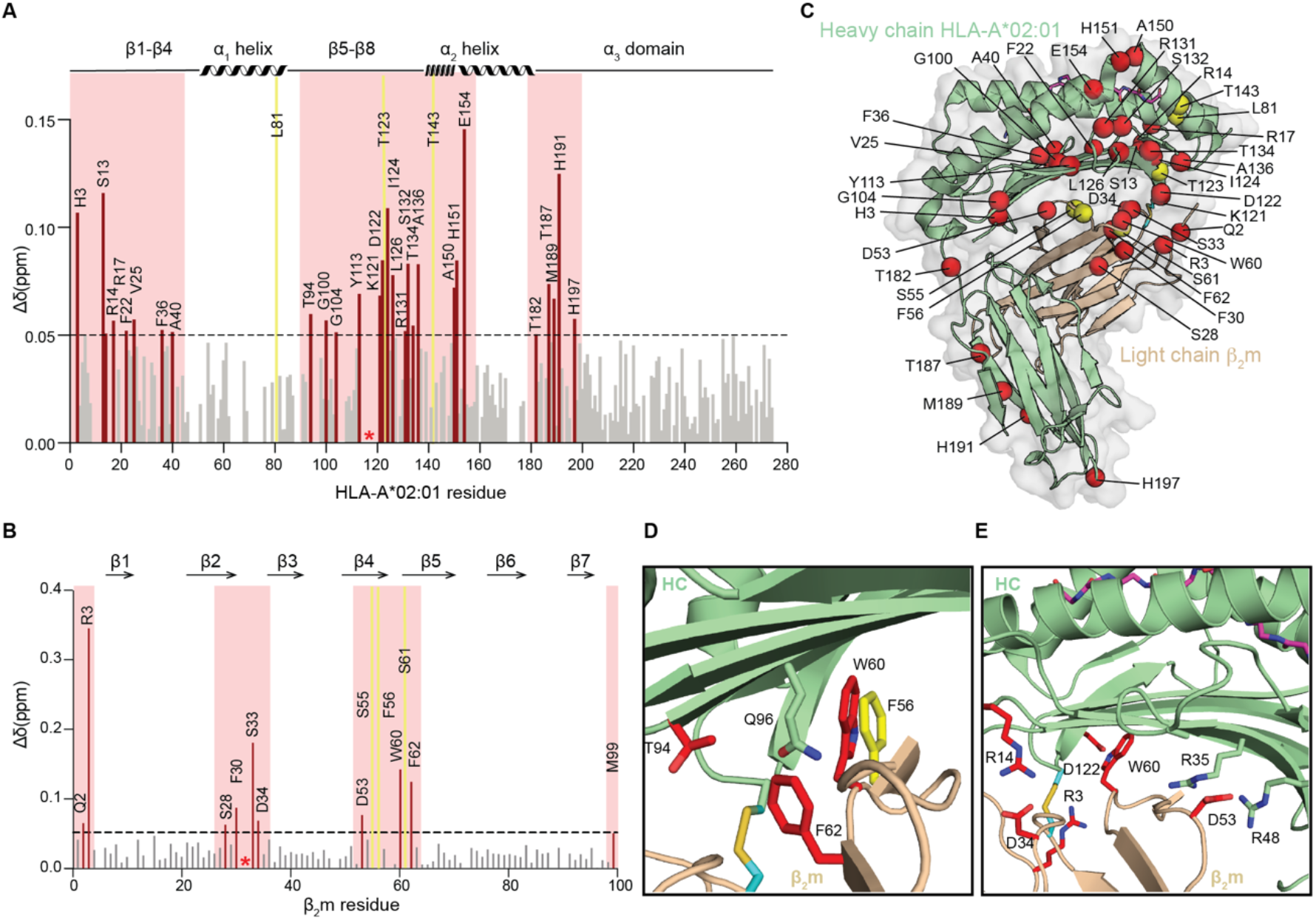
Disulfide-engineered pHLA-A*02:01 shows induced conformational adaptations in solution. **A-B.** Calculated CSPs between the WT and open HLA-A*02:01/β_2_m/MART1 are plotted as bar graphs across **A.** HC and **B.** β_2_m amide backbone. A significance threshold of 0.05 ppm is determined that is 5-fold higher than the ^1^H sensitivity of the NMR instrument. Residues with significant CSPs are highlighted in red, and exchange-broadened residues in the open HLA-A*02:01 relative to the WT are colored in yellow. Cysteine mutations (G120C and H31C) are indicated by a red asterisk. **C.** Residues with CSPs above the significance threshold and exchange broadened in the open HLA-A*02:01/β_2_m/MART1 are plotted as red and yellow spheres for the amine, respectively, on a representative HLA-A*02:01/β_2_m/MART1 crystal structure (PDB ID: 3MRQ). **D-E.** Enlarged images of **D.** hydrophobic residues near the disulfide bond and **E.** residues within the hydrogen network. Side chains are displayed and highlighted in red for significant CSPs and yellow for exchange-broadening.

In total, we identified 38 residues that were significantly affected by the formation of the interchain disulfide bond, signifying substantial, global differences between the conformational ensembles sampled by open and WT MHC-I molecules in solution (**Fig. 2A, B**). As expected, most of the impacted residues were found near the HC and β_2_m interface in the region surrounding the disulfide linkage (G120C and H31C) (**Fig. 2C**). Particularly, the β-sheet floor of the peptide binding groove, including the C, D, E, and F pockets showed high CSP values (**Fig. 2A, C**). Arginine at position 3, located on the flexible loop region close to the engineered disulfide bond, was the most affected residue on the β_2_m subunit (**Fig. 2B, C**). These effects indicate local structural rearrangements, induced at the vicinity of the disulfide linkage. Remarkably, our NMR data also indicate CSPs at residue W60 in β_2_m, while F56 displayed exchange broadening in the open but not in the WT, indicating altered microsecond to millisecond timescale dynamics (**Fig. 3D**). As shown in previous studies, the species-conserved F56 and W60 in β_2_m play a central role in stabilizing the interface with the α_1_α_2_ domain, acting as a conformational switch which controls peptide binding and release(35, 49, 50). Additionally, the HLA allele-conserved residues F8, T10, Q96, and M98 form the central part of a hydrophobic pocket together with F56, W60, and F62 from β_2_m(35). Therefore, the conformational changes observed for these residues upon covalently associating the HC and β_2_m may contribute to the overall stabilization of the peptide-loaded MHC-I, given the known role of β_2_m in promoting an allosteric enhancement of peptide binding(51,52).

**Figure 3.**
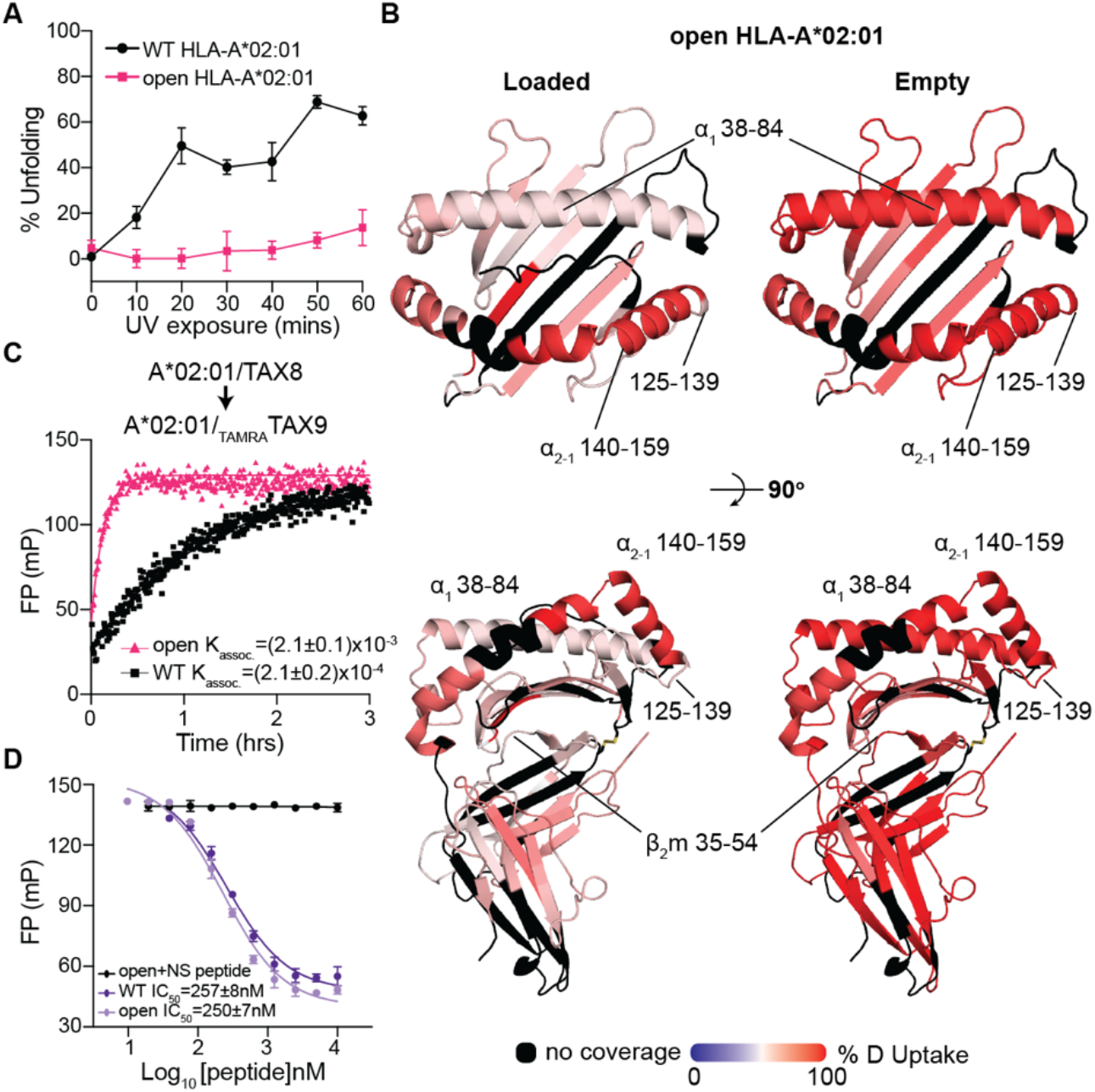
Engineered disulfide stabilizes MHC-I at an open, peptide-receptive conformation. **A.** Percent unfolding defined by the normalized fluorescent intensity at 25°C for the WT or open HLA-A*02:01/KILGFVFJV upon UV irradiation. The duration of UV irradiation is shown on the x-axis. Results of three technical replicates (mean ± σ) are plotted. **B.** Percent deuterium uptake resolved to individual residues upon 600-second deuterium labeling for peptide-loaded (left) and empty (right) states are mapped onto the HLA-A*02:01 crystal structure (PDB ID: 1DUZ) for visualization. Red and blue colored regions indicate segments containing peptides with 100% ΔHDX (red—more deuteration) or 0% ΔHDX (blue—less deuteration), respectively; black indicates regions where peptides were not obtained for peptide-loaded and empty protein states. **C.** Association profiles of the fluorophore-conjugated peptide _TAMRA_TAX9 to the WT or open HLA-A*02:01/TAX8, as indicated. Results of three replicates (mean) are plotted. **D.** Competitive binding of _TAMRA_TAX9 to the WT or open HLA-A*02:01/TAX8 as a function of increasing peptide concentration, measured by fluorescence polarization. An irrelevant peptide, p29 (YPNVNIHNF), was used as a negative control.

Moreover, residues H31 and W60 in β_2_m were also known to participate in a hydrogen bond network together with residues Q96, G120, and D122 in the α_1_α_2_ domains(35, 50). Our NMR data show that disulfide bond formation rearranges this network, including R3, D34, D53, and W60 on hβ_2_m and corresponding R14, R35, R48, and D122 on the HC (**Fig. 3E**). In addition, we observe long-range CSPs on the α_2-1_ helix and the far end of the β-sandwich fold on the α_3_ domain (**Fig. 2A, C**). This long-range effect supports our hypothesis that the interchain disulfide can trigger substantial global changes in protein dynamics, since the α_2-1_ helix has been previously shown to transition between open and closed states of the MHC-I groove for peptide loading. Similarly, our CSP data show a pronounced long-range effect suggesting a repacking of residues T187, M189, H191, and H197 within the α_3_ domain, via a lever arm effect transduced through residue T182 located on the loop joining the α_2_ and α_3_ domains. Thus, these results collectively demonstrate extensive local and long-range structural changes introduced by the bridging disulfide between the HC and β_2_m. Further, our NMR data suggest that the engineered disulfide bond may enhance peptide loading by inducing an allosteric conformational change of the peptide binding groove.

### Interchain disulfide bond formation induces a peptide-receptive MHC-I conformation

To test our hypothesis that the covalent linkage association between the β_2_m and α_1_α_2_ interface can improve the overall stability of empty MHC-I molecules, we used DSF to compare the percent unfolding of WT and open HLA-A*02:01 refolded with a photo-sensitive peptide upon varying periods of UV irradiation. While, in the absence of a rescuing peptide, WT molecules showed increasing amounts of protein denaturation, as measured by increased binding to the hydrophobic SYPRO orange, open heterodimers showed no substantial increase in the amount of unfolded protein leading to a 5-fold higher percent unfolding for the WT upon 1-hour UV irradiation (**Fig. 3B**). This is consistent with our previous DSF results showing that open molecules exhibit higher thermal stabilities when either empty or loaded with low-to moderate-affinity peptides (**Fig. 1D**). We next performed hydrogen-deuterium exchange-mass spectrometry (HDX-MS)(53) to identify differences in solvent accessibility patterns between open HLA-A*02:01 molecules in their peptide-loaded and empty states. Tandem analysis of the percent deuterium uptake as a function of exchange reaction time for different peptide fragments revealed exchange saturation within 600 seconds (**Fig. S4**). We observed significant differences in HDX patterns at specific regions, including the peptide binding groove, the α_3_ domain, and the β_2_m subunit (**Fig. 3D**). We also found low deuterium exchange on regions corresponding to the α_1_ helix and β-sheet floor of the peptide binding groove for the loaded molecules. Previous studies for WT HLA-A*02:01 have established that the α_2-1_ helix shows high deuterium uptake in the empty state(26, 54), likely due to an approximate 3 Å widening of the groove seen in crystal structures(13, 45, 55). In contrast, our HDX data recorded for open HLA-A*02:01 showed that residues 140-159 on the α_2-1_ helix have a similar level of deuterium uptake between the peptide-loaded and empty molecules (**Fig. 3B, Fig. S4**). These results suggest that bridging the HC/β_2_m interface facilitates the transition between open and closed states, enhancing the exchange of peptide ligands.

To examine whether the stabilizing effects of the MHC-I groove revealed by our NMR and HDX data bear any functional consequences in promoting peptide exchange, we compared the binding traces of a fluorophore-labeled peptide to HLA-A*02:01 molecules refolded with the suboptimal placeholder peptide TAX8 (56). The observed apparent association rates (Kassoc.) were determined by fitting a one-phase association model. We incubated the same concentration of refolded WT or open TAX8/HLA-A*02:01 complexes with a fluorescently labeled _TAMRA_TAX9 peptide, and measured a 10-fold higher exchange rate for the open molecules (**Fig. 3C**). Accordingly, high-affinity TAX9 could be readily loaded into the open or WT HLA-A*02:01 molecules, out-competing for the binding of fluorescent TAX9 (**Fig. 3D**). However, despite showing different peptide exchange kinetics, TAX9 showed identical IC_50_ values (approx. 200 nanomolar range) between WT and open HLA-A*02:01 (**Fig. 3C, D**). Taken together, these results support that the engineered disulfide bond indeed allosterically induces an open conformation of the MHC-I groove to enhance exchange kinetics, albeit without influencing the free energy of peptide binding.

### Open MHC-I molecules promote ligand exchange on a broad repertoire of HLA allotypes

We next sought to investigate how stabilizing the open MHC-I conformation may contribute to enhanced peptide exchange kinetics. To do this, we developed independent FP assays under conditions that allowed us to monitor the dissociation or association of _TAMRA_TAX9 to WT versus open HLA-A*02:01 molecules (**Fig. 4A, B**). When HLA-A*02:01 was refolded with the high-affinity _TAMRA_TAX9 probe and incubated with a large molar excess of competing TAX9 peptide, open molecules demonstrated a minor decrease in polarization anisotropy relative to the WT, indicating accelerated dissociation of _TAMRA_TAX9 from the MHC-I groove (**Fig. 4C**). This is due to the presence of the high-affinity (nanomolar range *K*_D_) _TAMRA_TAX9 peptide with a slow dissociation rate from both molecules, becoming the rate-limiting step in the overall reaction scheme (**Fig. 4A**). Conversely, when probing peptide association (**Fig. 4A**) open HLA-A*02:01 molecules refolded with the intermediate affinity placeholder peptide TAX8 showed a much higher rate of exchange with the _TAMRA_TAX9 probe (**Fig. 4C, Fig. S5**), reaching a higher plateau at steady-state. In agreement with our established peptide dissociation experiments, these results indicate noticeably higher amounts of stable, peptide-receptive molecules for the open variant than the WT at the same overall protein concentration. To quantitatively compare the effect of disulfide bridging on peptide exchange kinetics for different MHC-I systems, we measured an apparent association rate constant, k^app^_on_, defined as the slope of the linear correlation between K_assoc_. and the TAX8/HLA-A*02:01protein concentration. Under the first-order reaction scheme shown in Fig. 4B, this rate is proportional to the concentration of empty, receptive HLA-A*02:01 molecules in the system (**Fig. 4B, D**). Open HLA-A*02:01, compared to the WT, exhibited a more than 10-fold enhancement of the apparent k_on_, which is determined by both the formation of receptive molecules and the stability of these empty molecules (**Fig. 4D**). Taken together, open HLA-A*02:01 demonstrated faster kinetics of peptide exchange, likely because the rate-limiting transition, the intermediate step required to generate empty, receptive molecules, is faster due to the allosteric effects on α_2-1_ helix of the peptide binding groove originating from stable, covalent association with β_2_m, as shown by our NMR and HDX-MS data (**Fig 2, Fig. 3B**).

**Figure 4.**
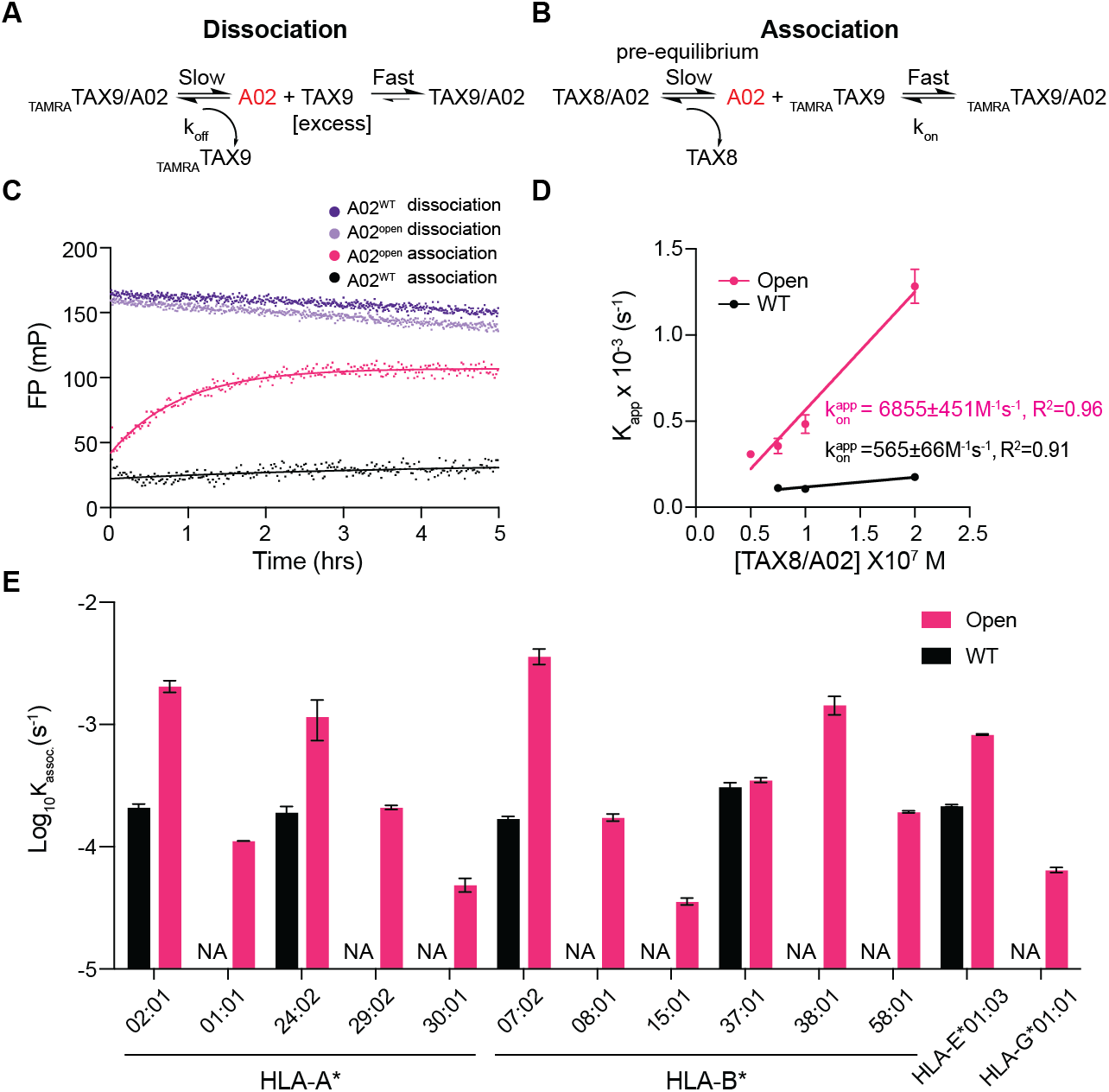
Open MHC-I improves peptide exchange efficiency on a broad repertoire of HLA allotypes. **A-B.** Schematic summary of fitted kinetics obtained from FP analyses of peptide exchange, showing **A.** the dissociation of 40 nM _TAMRA_TAX9/A02 in the presence of 1 uM unlabeled TAX9 and **B**. 40 nM _TAMRA_TAX9 in different concentrations of TAX8/A02 (50, 75, 100, and 200nM). **C.** The dissociation profiles of 40 nM WT or open _TAMRA_TAX9/A02 in the presence of 1 μM unlabeled TAX9, and association profiles of 40 nM WT or open _TAMRA_TAX9 in 50 nM TAX8/A02, as indicated. **D**. Linear correlations between the apparent rate constants K_assoc_. and the concentrations of TAX8/A02. The extrapolation of the slope between K_assoc_. and the concentrations of TAX8/A02 determine the apparent association rate kon. **E.** Log-scale comparison of K_assoc_. for the WT (black) or the open (pink) HLA-A*02:01, A*01:01, A*24:02, A*29:02, A*30:01, B*07:02, B*08:01, B*15:01, B*37:01, B*38:01, B*58:01, E*01:03, and G*01:01. The apparent rate constant K_assoc_. was determined by fitting the raw trace to a monoexponential association model. NA indicates no fitted K_assoc_.. Results of three technical replicates (mean ± σ) are plotted.

Empty, open MHC-I molecules can exist as a pre-equilibrium with the placeholder peptide-bound state to enable a rapid association with any high-affinity peptide ligand, in a ready-to-load manner. We further hypothesized that the interchain disulfide engineering could be applied to different HLA allotypes resulting in a universal, open MHC-I platform for antigen screening experiments. To quantitatively compare peptide exchange rates across different alleles, we then performed a series of FP experiments using optimized placeholder peptides, pHLA concentrations, and the protocol(26), where the binding of high-affinity fluorophore-labeled peptides was monitored through an increase in polarization (**Fig. S6**). Representatives covering five HLA-A and all HLA-B supertypes (A01, A02, A0103, A0124, A24 and B07, B08, B27, B44, B58, B62) were selected based on their global allelic frequency (**Table S2**). Additionally, we extended the study to cover the oligomorphic class Ib molecules, namely HLA-E*01:03 and HLA-G*01:01 (**Table S2**).

Our FP results showed that open MHC-I molecules (**Table S3**) demonstrate improved peptide exchange efficiency compared to the WT. Like open HLA-A*02:01 molecules, open HLA-B*07:02 exhibited a more than 20-fold increase in the apparent rate constant K_assoc_. (**Fig. 4E, Fig. S6E**). Both open HLA-A*24:02 and HLA-E*01:03 displayed enhanced peptide exchange kinetics by approximately 6- and 4-fold (**Fig. 4E, Fig. S6B, K**). The remaining allotypes, HLA-A*01:01, A*29:02, A*30:01, B*08:01, B*15:01, B*38:01, B*58:01, and G*01:01, showed a fitted K_assoc_. only in their open forms rather than in their WT counterparts (**Fig. 4E, Fig. S6**). Overall, we consistently observed a stabilizing effect on low to moderate-affinity peptide-loaded molecules across alleles (WT T_m_ < 53°C) (**Table. S4**). When loaded with a high-affinity peptide (WT T_m_ ≥ 53°C), T_m_ values generally stayed the same between the open and the WT, except for HLA-G*01:01 (**Table. S4**). Noticeably, suboptimally loaded HLA-B*37:01 in both open or WT format exhibited similar thermal stabilities and peptide exchange kinetics, revealing that receptive, empty molecules were preexisting in the sample for peptide binding. Although open MHC-I demonstrated fast exchange kinetics, we showed that two type 1 diabetes (T1D) epitopic peptides, HLVEALYLV and ALIDVFHQY, have the same IC_50_ towards both the WT and open variants encompassing HLA-A*02:01 and HLA-A*29:02 (**Fig. S7**). In summary, our FP results demonstrate that a wide range of open HLA allotypes exhibit enhanced thermal stabilities when loaded with suboptimal peptides, and greatly accelerated peptide exchange efficiency without compromising the stability of the resulting high-affinity pHLA complexes. These results provide additional evidence to support our hypothesis that the interchain disulfide bond offers an adaptable structural feature, which stabilizes a receptive MHC-I state, therefore enabling the spontaneous loading of peptide ligands across polymorphic HLA allotypes.

### Application of open MHC-I as molecular probes for T cell detection and ligand screening

We finally evaluated the use of open MHC-I molecules as ready-to-load reagents in tetramer-based T cell detection strategies. We conducted 1-hour peptide exchange reactions at room temperature (RT) for both WT and open HLA-A*02:01 loaded with a placeholder peptide (KILGIVFβFV(26)) for an established tumor-associated antigen, NY-ESO-1 (SLLMWITQV). BSP-tagged HLA-A*02:01 were purified, biotinylated, and tetramerized using streptavidin labeled with predefined fluorochromes (**Fig. 5A, Fig. S8**). We then stained primary CD8+ T cells transduced with the TCR 1G4 that recognizes the NY-ESO-1 peptide displayed by HLA-A*02:01(57). Compared to WT HLA-A*02:01/NY-ESO-1, tetramers generated with open molecules exhibited a similar staining level (**Fig. 5B**). We used non-exchanged HLA-A*02:01/KILGIVFβFV molecules as negative controls. Analysis by flow cytometry showed minimal levels of background staining using the open tetramers (**Fig. 5C**), showing that the 1G4 was not able to recognize HLA-A*02:01/KILGIVFβFV. However, the WT tetramers demonstrated a higher level of background staining, exhibiting one order of magnitude lower intensity relative to the WT HLA-A*02:01/NY-ESO-1 tetramer staining levels (**Fig. 5C**), likely due to the formation of empty MHC-I heterodimers, which can interact with the CD8 co-receptor(58). The noticeable difference in background staining might also be caused by varying levels of formation of protein aggregates induced by peptide dissociation and subsequent loss of β_2_m. Despite having rapid peptide exchange kinetics, disulfide-engineered open MHC-I demonstrated comparable IC_50_ values for binding of high-affinity epitopes to the WT and heterodimer stabilization in a conformation that is receptive to peptides (**Fig. 3D, Fig. S7**). In addition, open HLA-A*02:01 loaded with moderate-affinity peptides, SLLMWITQC and SLLMWITQA (NYESO C and A), via exchange reaction show consistent T cell staining (**Fig. 5D**). Thus, the engineered disulfide does not interfere with the peptide binding and interactions with T cell receptors. Having a reduced level of background staining allows the use of higher concentrations of tetramers to study interactions with low-affinity TCRs, as seen, for example, in the case of autoimmune peptide epitopes(64). Using open MHC-I as a ready-to-load system can help elucidate the intrinsic peptide selector function across different alleles to optimize peptide binding motifs, but also has important ramifications for developing combinatorial barcoded libraries of pHLA antigens toward TCR repertoire characterization(59).

**Figure 5.**
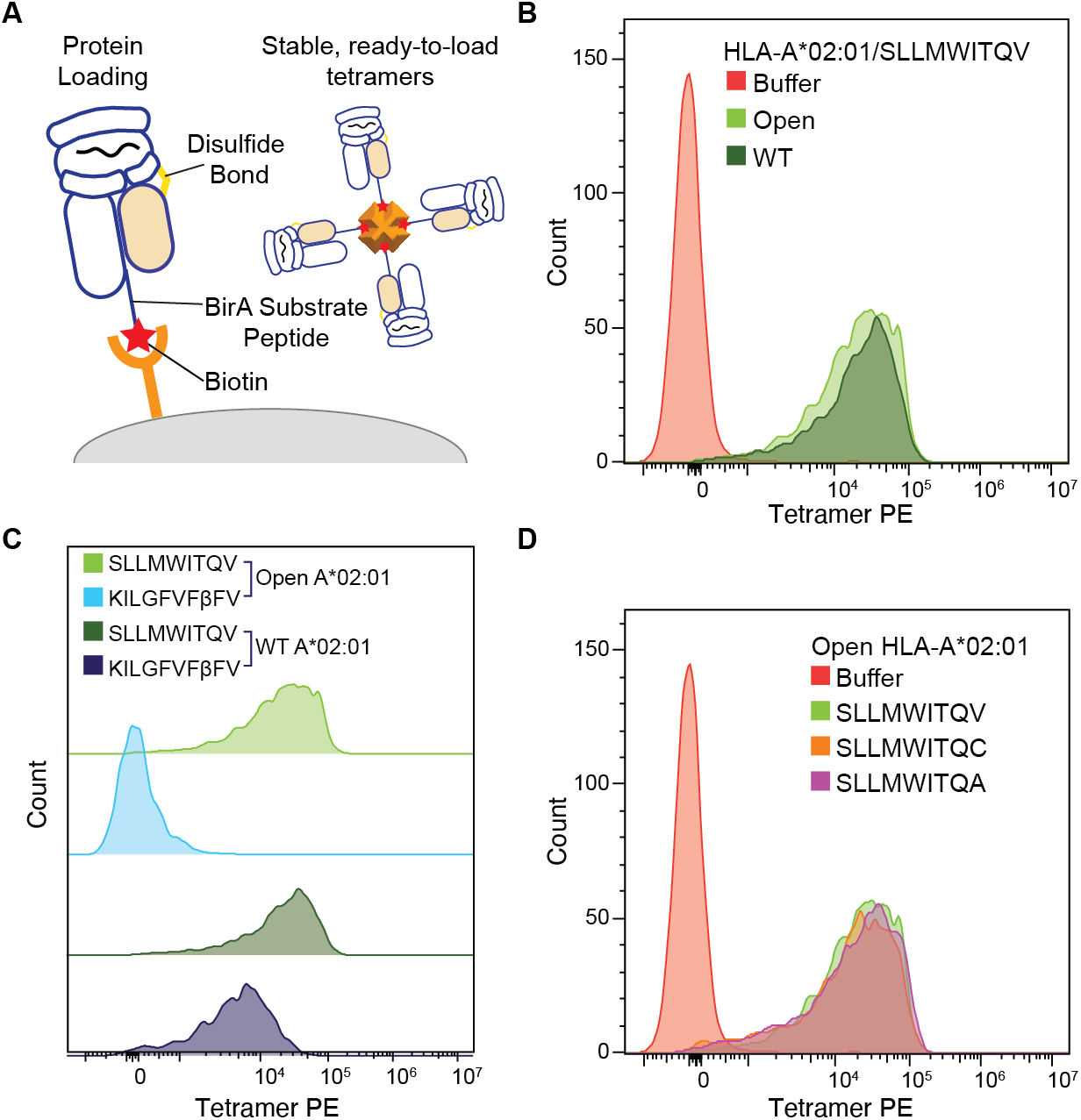
Open HLA-A*02:01 molecules enable effective T cell detection by reducing non-specific background staining compared to the WT. **A.** A schematic summary of the disulfide-linked HLA-I molecules with the desired BSA tag enables biotinylation and sets the stage for tetramerization. **B.** Staining of 1G4-transduced primary CD8+ T cells with PE-tetramers of open and WT HLA-A*02:01/NY-ESO-1(V), light and dark green, respectively. **C.** Staining of 1G4-transduced primary CD8+ T cells with PE-tetramers of open and WT HLA-A*02:01/NY-ESO-1(V), light and dark green, compared to open and WT HLA-A*02:01 loaded with a non-specific peptide, light and dark blue. **D.** Staining of 1 G4-transduced primary CD8+ T cells with PE-tetramers of open HLA-A*02:01 loaded with different NY-ESO-1 peptides SLLMWITQV (light green), SLLMWITQVC (orange), and SLLMWITQA (purple).

Finally, we extended the design of open HLA-I to nonclassical MHC-I, MR1, which can present small molecule metabolites via both non-covalent and covalent loading by the A pocket (**Fig. S9A**). We demonstrated that the open MR1 C262S could be refolded in vitro with covalently linked molecules Ac-6-FP and the non-covalent molecule DCF with a noticeable improvement in protein yield (**Fig. S9B, C**). Consistently, we observed a substantial upwards shift of the T_m_ by more than 10°C for the open MR1/Ac-6-FP than the WT (**Fig. S9D**). HLA-F*01:01 molecules known to be stable in their empty form and accommodating long peptides with a length range from 7 to >30 amino acids averaging at 12 residues(60) can also adapt the same structural design to generate open heterodimers (**Fig. S9E**). These results further support the universality of the open platform, covering not only classical HLA Ia allotypes but also the oligomorphic HLA Ib and nonclassical MHC-I.

## Discussion

The inherent instability of peptide-deficient MHC-I heterodimers is a major pitfall of current peptide exchange technologies, limiting screening applications for important therapeutic antigens. Our combined biochemical and biophysical characterization outlines a universal design strategy for generating ready-to-load MHC-I conformers across various disease-relevant HLA allotypes. Compared to the UV- or heat-induced peptide exchange methods(19, 21), open MHC-I molecules combined with β-peptide “goldilocks” ligands introduce mild exchange conditions suitable for large-scale screening applications. Previous work has highlighted the potential of chaperone-mediated exchange in various settings (24, 61, 62). While tapasin has shown preferential binding to HLA-B alleles, TAPBPR preferably interacts with HLA-A alleles but mainly covers the A02 and A24 supertypes(46, 63). More recent work has expanded the TAPBPR-mediated peptide exchange on a broad repertoire of allotypes using TAPBPR orthologs and engineered variants(26). However, compared to the open MHC-I platform, the approach requires optimized placeholder peptides and recombinant chaperone proteins to stabilize empty, receptive molecules. Ready-to-load MHC-I molecules have been derived through the introduction of an engineered disulfide bond between WT residues Tyr 84 and Ala 139, linking the α_1_ and α_2_ helices at the F pocket to stabilize MHC-I molecules in an empty, receptive conformation(32). However, this approach has been applied in limited HLA alleles(30–34). Open MHC-I molecules exploit the positive cooperativity between peptide association and β_2_m binding to the HC(16, 38) to stabilize the peptide binding groove in an open conformation without directly altering the properties of the MHC-I peptide binding groove. Instead, our approach exploits the known allosteric switch connecting W60 from β_2_m with the floor of the MHC-I groove and α_2-1_ helix, to generate molecules with favorable exchange properties. We show the generality of the design by aligning both sequences and structures and demonstrating peptide exchange applications for representatives from five HLA-A and all HLA-B supertypes.

Our complementary FP assays show that both WT and open MHC-I molecules loaded with moderate-affinity placeholder peptides undergo a slow but spontaneous peptide unloading process to generate empty molecules (**Fig. 6A**). While such empty WT MHC-I heterodimers are intrinsically unstable and susceptible to β_2_m loss and irreversible heavy chain aggregation through the exposure of hydrophobic surfaces, open MHC-I molecules maintain a soluble reservoir of receptive molecules for peptide loading (**Fig. 6B**). Previous studies have demonstrated that the flexibility of the α_2-1_ helix allows the peptide binding groove to dynamically shift between an open and closed state for peptide exchange(6, 64). Binding to high-affinity peptides triggers the closed conformation, and promotes the dissociation of molecular chaperones, which are known to recognize an open conformational epitope at the α_2-1_ helix(13, 46, 65, 66). Using solution NMR, we have demonstrated that our open MHC-I molecules undergo an allosteric conformational change of the α_2-1_ helix, which enables the rapid capture of incoming peptides without perturbing the global stability of the resulting pMHC-I product. Open MHC-I molecules, therefore, enhance the rate of generating receptive molecules, which is the rate-limiting step in the overall peptide exchange reaction scheme. Our HDX-MS and DSF data reveal an increase in solvent exposure for the α_2-1_ helix in both peptide-loaded and empty states without compromising the thermal stabilities of open MHC-I molecules. This indicates that the energy barrier separating the open and closed states might be minimized, sequentially lowering the activation free energy for peptide unloading (**Fig. 6C**). Therefore, the covalently linked β_2_m further functions as a conformational chaperone to allosterically induce the open conformation of the peptide binding groove, resulting in rapid peptide loading and unloading in favor of high-affinity over placeholder peptides.

**Figure 6.**
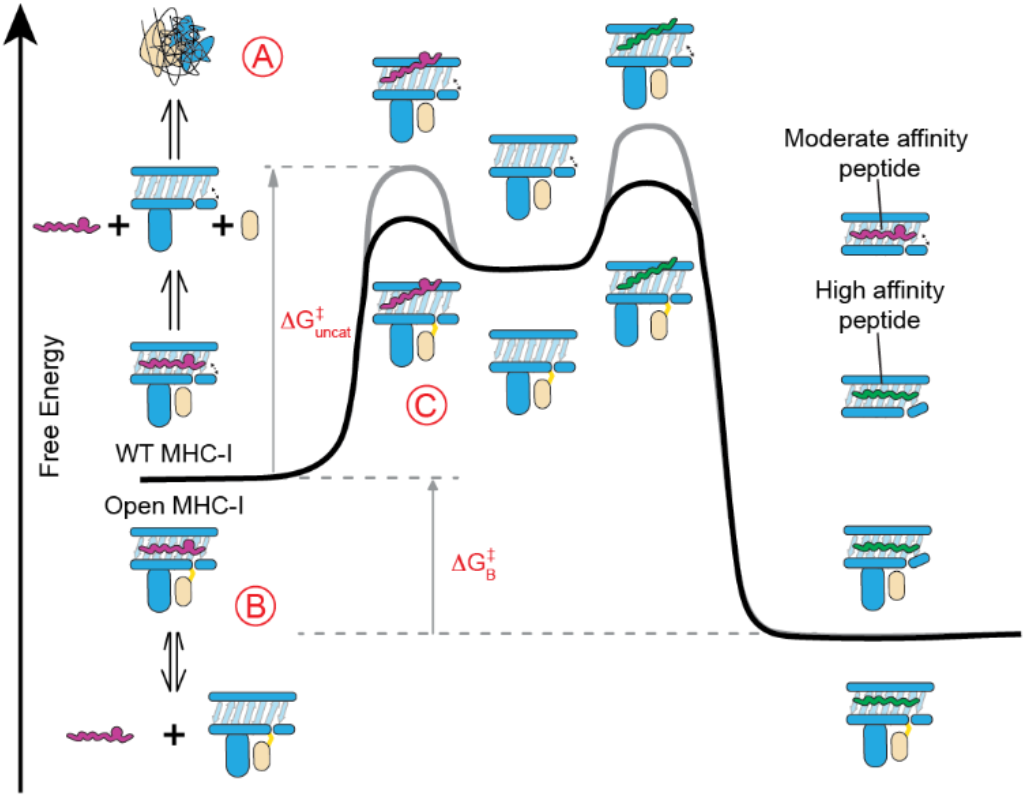
Open MHC-I molecules modulate the free energy landscape of peptide exchange. **A.** WT MHC-I molecules loaded with moderate-affinity placeholder peptides can spontaneously generate empty molecules, leading to β_2_m loss and irreversible protein aggregation. **B-C.** Open MHC-I enhances peptide exchange by **B.** stabilizing empty molecules to prevent their aggregation and **C.** lowering the activation free energy (ΔG_uncat_) for peptide un-loading via a stabilized open conformation.

Open MHC-I allows for minimal protein modifications leading to enhanced exchange reactions across allotypes. Thus, these molecules could be a versatile tool for screening antigenic epitopes, enabling the detection of low-frequency receptors. Further, it is necessary to confirm that open MHC-I molecules are not interfering with peptide repertoire selection or interactions with cognate TCRs. A more detailed study of thermodynamic parameters relevant to peptide binding using Isothermal Titration Calorimetry (ITC) will provide additional insights into how the presence of the interchain disulfide modulates the peptide-free energy landscape. Additionally, a follow-up study using the open MHC-I focusing on detecting antigen-specific T cells across different HLA allotypes is required to demonstrate the broad usage of this platform as an off-the-shelf reagent. In summary, we outline an alternative structure-guided design of open MHC-I molecules that are conformationally stable and ligand-receptive across five HLA-A and all HLA-B supertypes, oligomorphic HLA-Ib alleles, HLA-E, -G, and -F, and nonclassical MHC-like molecules, MR1. Our data provide a framework for exploring the allosteric networks that exist in the structures of native MHC-I molecules to further guide the design of ultra-stabilized, universal ligand exchange technologies, which can be used to address highly polymorphic HLA allotypes.

## Acknowledgments

This work was supported through grants by NIAID (5R01AI143997), NIDDK (5U01DK112217), and NIGMS (5R35GM125034) to N.G.S. and NIGMS (R35GM142505) to G.M.B. We acknowledge NIH training grant (T32GM132039) for support to C.H.W. We acknowledge Dr. Andy J. Minn and Devin Dersh (University of Pennsylvania) for providing the primary CD8+ cell lines for cell staining and flow cytometry. We acknowledge Nick Marotta for performing MR1 protein refolding experiments, Andrew C. McShan and Tyler J. Florio for assistance using the Disulfide by Design web server, Ananya Majumdar for assistance with data collection on the 800 MHz spectrometer at Johns Hopkins University, Leland Mayne for advising the HDX-MS experiments, and the Human Immunology Core at the University of Pennsylvania providing all primary cells used in this study.

## Data Availability

NMR assignments for the wild type and open A*02/MART1 complexes (for both the heavy and hβ_2_m light chains) have been deposited into the Biological Magnetic Resonance Data Bank (http://www.bmrb.wisc.edu) under accession numbers 51101 and 51781 respectively.

