## Supplementary Methods, Figures and Tables for "Universal open MHC-I molecules for rapid peptide loading and enhanced complex stability across HLA allotypes"

**This PDF file includes:**

Supplemental Methods
Supplementary Figures S1 to S9
Tables S1 to S4

### Supplemental Methods

**Peptides and ligands**. All peptide sequences are given as standard single-letter codes. Peptides for different HLA allotypes were selected by NetMHCpan4.1 and purchased from Genscript, Piscataway, USA, at >90% purity. L-β-Phenylalanine (βF) containing placeholder peptides were synthesized in-house on 2-chlorotrityl resin using a CEM Liberty Blue automated microwave peptide synthesizer from Fmoc protected amino acids (including Fmoc-β-Phe-OH) employing iterative cycles of N, N'-Diisopropylcarbodiimide (DIC)/Ethyl cyanohydroxyiminoacetate (Oxyma) mediated coupling and piperidine mediated deprotection, both under microwave irradiation. Peptides were deprotected and cleaved from the resin by treatment with trifluoroacetic acid/water/triisoproylsilane/phenol (88:5:5:2) for 1-3 hours. The solvent was removed under a flow of nitrogen, and peptides were precipitated with ice-cold ether. Peptides were subsequently purified by reverse phase chromatography eluting with 5-95% acetonitrile in water containing 0.05% trifluoroacetic acid over a C18 column. Peaks containing peptides were identified by LC-MS, pooled, and concentrated in vacuo to yield a colorless solid. Photosensitive peptides were purchased from Biopeptek Inc, Malvern, USA, or synthesized in-house using Fmoc-3-amino-3-(2-nitrophenyl)-propionic acid (J). Peptides were solubilized in distilled water and centrifuged at 14000 rpm for 15 minutes. The concentration of each peptide solution was measured and calculated using the respective absorbance and extinction coefficient at 205 nm wavelength. MR1 C262S ligand acetyl-6-formylpterin (Ac-6-FP) and diclofenac (DCF) were purchased from Cayman Chemical (#23303) and Sigma D6899-10G.

**Recombinant protein expression, refolding, and purification**. Plasmid DNA encoding the BirA Substrate Peptide (BSP, LHHILDAQKMVWNHR)-tagged luminal domain of MHC-I heavy chains and human $\beta_2$m were provided by the NIH tetramer facility (Emory University) and transformed into *Escherichia coli* BL21(DE3) cells (New England Biolabs). Open heavy chains (G120C) and $\beta_2$m (H31C) were generated using site-directed mutagenesis and transformed into *Escherichia coli* BL21(DE3) cells using the pET-22b(+) vector. Cells were grown and harvested in the Luria-Broth medium, and inclusion bodies were pelleted and purified as previously described(1). For the generation of pMHC-I molecules, *in vitro* refolding was performed by slowly diluting a 100 mg mixture of either wild type (WT) or open MHC-I and $\beta_2$m at a 1:3 molar ratio over 4 hours in refolding buffer (0.4 M L-Arginine HCl, 100 mM Tris pH 8, 2 mM EDTA, 5 mM reduced L-glutathione, 0.5 mM oxidized L-glutathione) supplemented with 10 mg of the peptide. The mixture was protected from light when refolded with photosensitive peptides. Refolding proceeded for 4 days, and proteins were purified by size exclusion chromatography (SEC) using a HiLoad 16/600 Superdex 75 pg column at 1 mL/min with 150 mM NaCl, 20 mM Tris buffer, pH 8.0. Purified proteins were further confirmed in reduced and non-reduced conditions using sodium dodecyl sulfate-polyacrylamide (SDS-PAGE) gel electrophoresis. MR1 refolding was performed by diluting a 90 mg mixture of either WT or open HC and $\beta_2$m at a 1:1.3 molar ratio overnight in refolding buffer supplemented with 5 mg of DCF or Ac-6-FP. Protein purification was performed as described above.

**Differential Scanning Fluorimetry**. Differential Scanning Fluorimetry (DSF) was used to assess the thermal stabilities of the WT and the open pMHC-I protein complexes. 7 µM of placeholder peptide-loaded MHC-I molecules were incubated with the desired peptide at a 1:10 molar ratio at room temperature (RT) overnight and then mixed with 10X SYPRO Orange dye in PBS buffer (150 mM NaCl, 20 mM sodium phosphate, pH 7.2) to a final volume of 20 µL. Samples were loaded into MicroAmp Optical 384 well plate and ran in triplicates. The experiment was performed on a QuantStudio™ 5 Real-Time PCR machine with excitation and emission wavelengths set to 470 nm and 569 nm. The temperature was incrementally increased at a rate of 1°C per minute between 25°C and 95°C. Data analysis and fitting were performed in GraphPad Prism v9. To determine the percent unfolding, WT and open HLA-A*02:01/KILGFVFJV were UV irradiated for 0, 10, 20, 30, 40, 50, and 60 minutes. The full DSF traces were recorded at a constant rate of 1°C per minute between 25°C and 95°C. The fluorescence intensity (I) at 25°C was then normalized against the maximum I. Data analysis and fitting were performed in GraphPad Prism v9.

**NMR sample preparation and methyl resonance assignment**. NMR samples of WT and open HLA-A*02:01/$\beta_2$m/MART1 complex were prepared with an [$^{15}$N, $^{13}$C, $^{2}$H] isotope-selective labeling scheme using established protocols and reagents(2, 3). The HC and $\beta_2$m components were each isotopically labeled independently using M9 media in *E. coli*(13) and refolded with the complementary complex components expressed at natural isotopic abundance, as described previously for the same system, to generate two NMR samples each for open and WT. Samples in the concentration range of 50 to 150 µM were prepared in a standard NMR buffer (150 mM NaCl, 20 mM sodium phosphate pH 7.2, 0.001 M sodium azide, 5% $D_2O$) in the presence of 2-fold molar excess of MART1 peptide, and all datasets were collected at 298-300 K. Backbone resonance assignments for the WT complexes were derived using a series of TROSY-based 2D and 3D experiments recorded at a $^{1}$H field of 600 or 800 MHz following a multi-pronged approach described previously for a similar system(4), including HNCO, HNCA, and HN(CA)CB triple-resonance experiments and SOFAST-based Hn-NHn NOESY experiments recorded at 800 MHz(5–9). Assignments were then transferred to the spectra of the open complexes and confirmed by TROSY-readout triple-resonance experiments (HNCO, HNCA, and HN(CA)CB), recorded at 600 MHz. Final backbone assignments were verified using the TALOS-N server (10) and deposited in the Biological Magnetic Resonance Bank (IDs: 51101 and 51781). For chemical shift perturbation calculations, the WT and open TROSY NMR spectra were aligned to each other using a residue far from the mutation sites as a reference based on an existing crystal structure, in a region where open and WT peaks were perfectly overlapped (HLA-A*02:01_E254 and $\beta_2$m_D96; PDB ID: 3mrq). Amide backbone chemical shift perturbations between the WT and the open variant were calculated using the following equation, given the aligned $^{15}$N and $^{1}$H chemical shifts: $\Delta\delta(ppm) = \sqrt{(\Delta\delta_H)^2 + \left(\frac{\Delta\delta_N}{10}\right)^2}$. All NMR data were processed with NMRPipe and analyzed using NMRFAM-SPARKY and POKY(11, 12).

**Hydrogen/Deuterium exchange mass spectrometry**. The open HLA-A*02:01/KILGFVFJV was dialyzed into equilibration buffer (150 mM NaCl, 20 mM sodium phosphate, pH 6.5 in $H_2O$) and diluted to a stock concentration of 30 μM and then either i) kept on ice without exposure to UV light or ii) UV-exposed for 45 min at 4°C. Samples were prepared and injected manually for several deuterium-exchange incubation periods. 5 μL open HLA-A*02:01/KILGFVFJV (30 μM) with or without UV-irradiation were diluted with 20 μL equilibration buffer (all H experiments, 0 s) or deuterium buffer (150 mM NaCl, 20 mM sodium phosphate pD 6.5 in $D_2O$) to 6 μM. The proteins were incubated with deuterium buffer for 20, 180, and 600 seconds at RT, and 15 minutes at 43°C for HLA-A*02:01/KILGFVFJV or at 34°C for UV-irradiated HLA-A*02:01/KILGFVFJV as all the D samples to calculate $\Delta Mass_{100\%}$. The samples were then quenched with an equal volume of acidic buffer (150 mM NaCl, 1 M TCEP, 20 mM sodium phosphate pH 2.35 in $H_2O$, 25 μL). Quenched proteins were immediately injected for LC-MS/MS in which integrated pepsin digestion was performed using a C8 5 μM column and a Q Exactive Orbitrap Mass Spectrometer. Peptide fragments corresponding to HLA- HLA-A*02:01 and $\beta_2$m were identified using Thermo Proteome Discoverer v2.4. The percent deuterium uptake was back-exchange corrected for each time point using the following equation(13): %D= $\frac{\Delta \mathrm{Mass}_T - \Delta \mathrm{Mass}_{0\%})}{\Delta \mathrm{Mass}_{100\%} - \Delta \mathrm{Mass}_{0\%})}$. ExMs2 program was used to identify and analyze deuterated peptides. The kinetic plots and the scaled B factor for the structure plot were generated by python3 and PyMOL(14).

**Fluorescence polarization**. The kinetic association of fluorescently labeled peptides and various peptide-loaded MHC-I was monitored by fluorescence polarization (FP). An optimized concentration of a fluorophore-labeled peptide (determined via serial dilution that yields a polarization baseline between 0 and 50 mP) was solubilized in FP buffer (150 mM NaCl, 20 mM sodium phosphate, 0.05% Tween-20, pH 7.4). MHC-I proteins and fluorophore-labeled peptides were directly added to the plate to 100 μL per well to avoid extended incubation and loss of data. The kinetic association was monitored for 2-12 hours, and polarization measurements were recorded every 28-105 seconds. The WT or open pMHC-I concentration remained constant across experiments at 200 nM, except for the MHC titration assays. Excitation and emission values used to monitor the fluorescence of TAMRA-labeled peptides were 531 and 595 nm, and FITC-labeled peptides were 475 and 525 nm. All experiments were performed in triplicates at RT. For $IC_{50}$ competition assays, a serial dilution of competitor peptide was added to 200 nM WT or open pMHC-I and the optimal concentration of fluorophore-labeled peptide. Kinetic association measurements were collected. Non-linear regression fitting allowed calculating plateau polarization (mP) values for each kinetic curve. Log transformed values of each peptide concentration were plotted against the plateau mP value, and an $IC_{50}$ curve was fit. Raw parallel ($I_{\parallel}$) and perpendicular emission intensities ($I_{\perp}$) were collected and converted to polarization (mP) values using the equation $1000*[(I_{\parallel}-(G*I_{\perp}))/(I_{\parallel}+(G*I_{\perp}))]$. An optimized G-factor was determined to be 0.33 for TAMRA-labeled peptides and 0.4 for FITC-labeled peptides in calculating baseline fluorescence and overall FP. The data analysis method was adapted and data fitting was performed in GraphPad Prism v9(15).

**Biotinylation and tetramer formation**. Biotinylation and tetramer formation of the WT and open HLA-A*02:01/KILGFVFβFV proteins were performed as previously described(16). The BSP-tagged proteins were biotinylated using the BirA biotin-protein ligase bulk reaction kit (Avidity), according to the manufacturer's instructions. Biotinylated molecules were washed using Amicon Ultra centrifugal filter units with a 100 kDa membrane cut-off, and the level of biotinylation was evaluated by SDS-PAGE gel shift assay in the presence of excess streptavidin. Biotinylated WT and open HLA-A*02:01/KILGFVFβFV were mixed with 10 fold molar excess of the NYESO-1 peptide variants, SLLMWITQV, SLLMWITQC, and SLLMWITQA. Each reaction was incubated 2 hours at room temperature and the peptide exchange reactions were confirmed by DSF. NYESO-1 peptide-loaded HLA-A*02:01 molecules were prepared at a final concentration of 2 mg/mL. Streptavidin-PE (Agilent Technologies, Inc.) at 4:1 monomer/streptavidin molar ratio was added to pMHC-I/$\beta_2$m over 10-time intervals every 10 mins at RT in the dark. The resulting pMHC-I/$\beta_2$m tetramers can be stored at 4°C for up to 4 weeks.

**1G4 TCR lentivirus production**. Lenti-X 293T cells (Takara) were cultured in DMEM (Gibco), 10% FBS (Gibco), and Glutamax (Gibco) and were plated one day before transfection. Cells were transfected at a confluency of 80-90% with TransIT-293 (Mirus) using pMD2.G (Addgene #12259, gift from Didier Trono), psPAX2 (Addgene #12260, gift from Didier Trono), and pSFFV-1G4. Virus-containing media was collected 24- and 48-hours post-transfection, clarified by centrifugation at 500 g for 10 min, and incubated with Lenti-X concentrator (Takara) for at least 24 hours. Virus was pooled and concentrated 50-100x, resuspended in PBS, aliquoted, and stored at -80°C for subsequent T cell infections.

**Primary human T cell tetramer staining**. The studies involving human participants were reviewed and approved by the University of Pennsylvania review board. Written informed consent to participate in this study was provided by the participants. Healthy donor T cells were processed by the Human Immunology Core at the University of Pennsylvania by magnetic separation of CD8+ T cells. Cells were cultured in Advanced RPMI (Gibco), 10% heat inactivated FBS (Gibco), Glutamax (Gibco), penicillin/streptomycin (Gibco), and 10mM HEPES (Quality Biological), supplemented with 300 U/mL recombinant IL-2 (NCI Biological Resources Branch). T cells were maintained at ~1 million cells/mL and were activated with a 1:1 ratio of Dynabeads Human T-Activator CD3/CD28 beads (Gibco) for 48 hours. 24 hours after initial activation, cells were either left untransduced or transduced with lentivirus expressing the 1G4 TCR. Cells were debeaded by magnetic separation and expanded in the presence of IL-2. Transduction efficiency was determined by staining with an anti-Vβ13.1-APC antibody (Miltenyi Biotec.), typically greater than 50%. Cells were cryopreserved with CryoStor CS10 (StemCell Technologies). Thawed T cells were recovered and regrown in IL-2-containing complete media for ~3 days prior to staining. Cells were harvested and washed with PBS/1% BSA/2 mM EDTA with 5 μg/mL PE-conjugated tetramer and incubated for 25 min at room temperature with slight shaking. After two washes with an RPMI-based wash buffer containing 1%

FBS, cells were resuspended in 1:1000 Sytox Blue diluted in wash buffer to distinguish dead cells. Samples were processed on an CytoFLEX LX and the data analyzed by FlowJo v10.8.1.

### Supplemental Figures

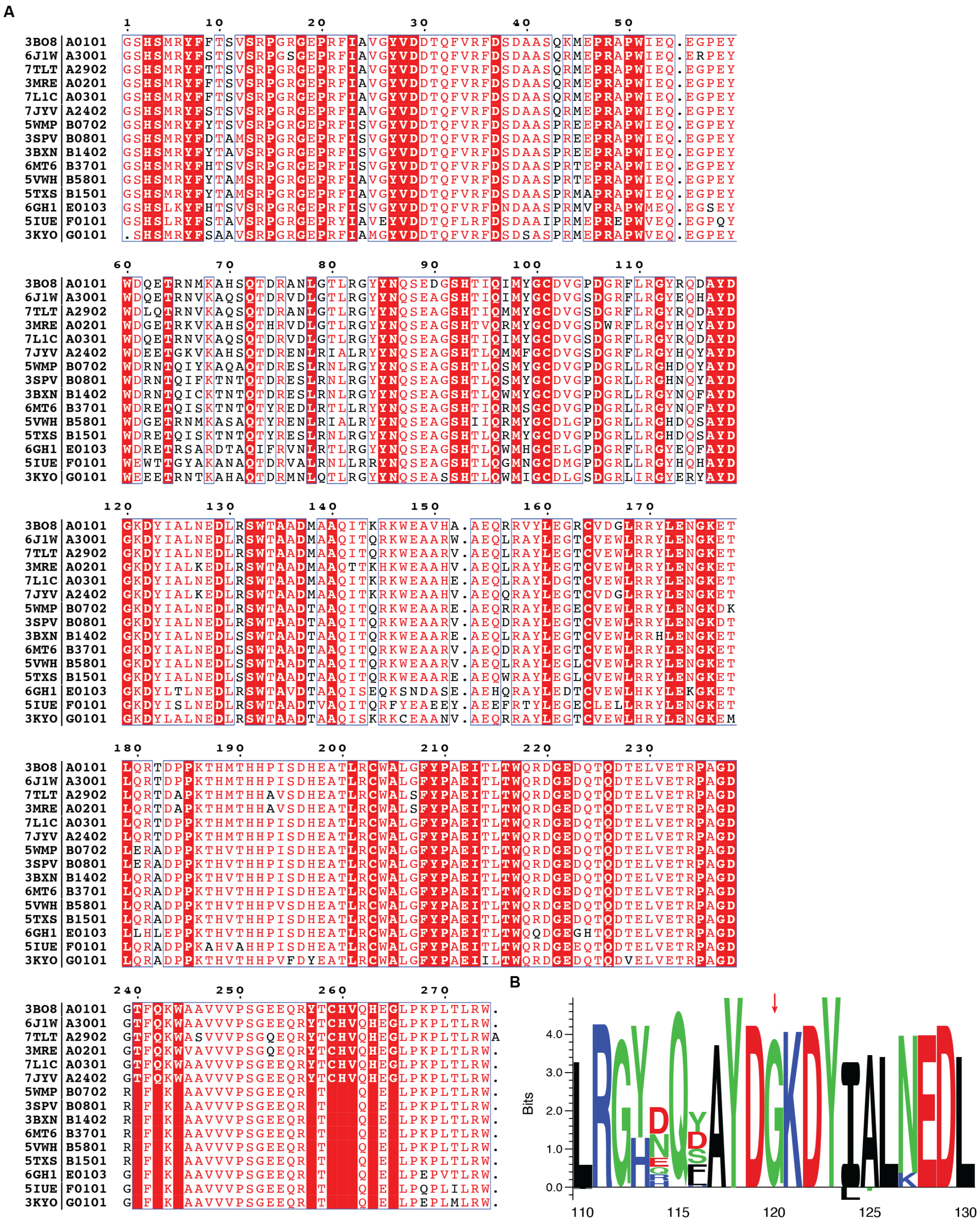


**Figure S1. Sequence alignment of distinct HLA allotypes across HLA-A, B, and C as well as HLA-Ib. A.** Sequence alignment of the HLA representatives were extracted from the Protein data bank and processed using ESPript(17), covering HLA-A and HLA-B supertypes as well as HLA-Ib, PDB ID as indicated. **B.** Seq2logo visualization(18) of the sequence alignment for 75 distinct HLA allotypes with >1% global population frequency shows a conserved residue G120. Sequence weighting used clustering, pseudo count with a weight of 0, and Kullback–Leibler logotype. The percentage frequency of amino acids on a specific position higher than 10% is shown on the positive *y*-axis, and less than 10% amino acids on the negative y-axis. Allele sequences were derived from the IPD-IMGT/HLA(19) and the alignment was performed using ClustalOmega(20).

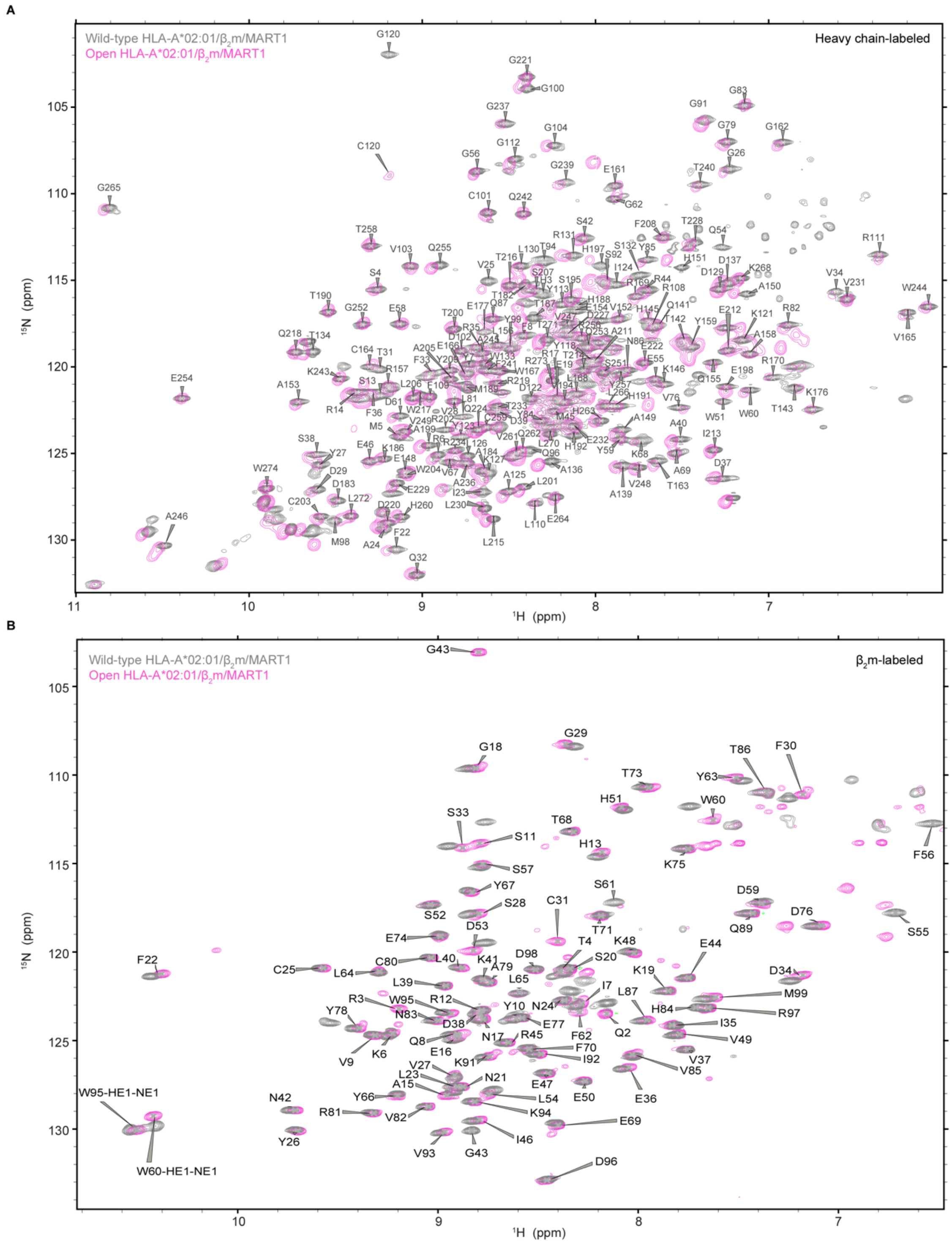


**Figure S2. Overlay of the WT and open MHC-I NMR spectra reveal substantial backbone chemical shift changes.** 2D $^{1}$H-$^{15}$N TROSY spectra of [$^{1}$H, $^{13}$C, $^{15}$N]-labeled **A.** HC (HLA-A*02:01) refolded with unlabeled light chain (β2m) and MART1 (ELAGIGILTV), or **B.** β2m bound to unlabeled HC and MART1. Spectra represent the WT complex collected at 800 MHz $^{1}$H magnetic field (in gray), overlayed by the open complex spectra collected at 600 MHz $^1$H magnetic field (in pink). All data were collected with identical buffer conditions (20 mM sodium phosphate, pH 7.2, and 150 mM NaCl) and at RT (298-300 K).

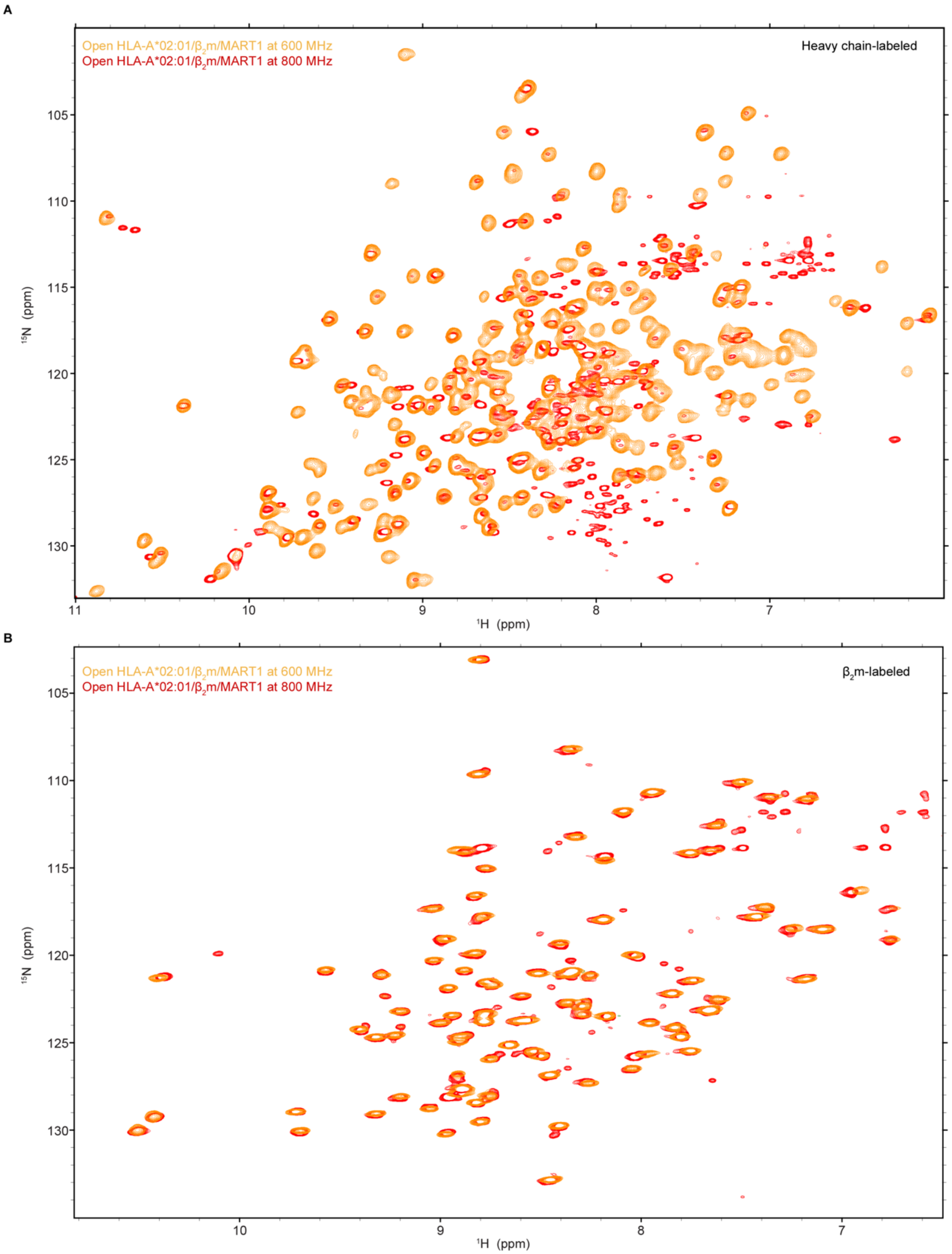


**Figure S3. Overlay of NMR spectra for the open MHC-I confirms the same backbone chemical shifts regardless of the magnetic field.** $^{1}$H-$^{15}$N TROSY data collected for the open HLA-A*02:01/$\beta_2$m/ MART1 at both 600 MHz (orange) and 800 MHz (red) for [$^{1}$H, $^{13}$C, $^{15}$N]-labeled HC refolded with $\beta_2$m and MART1 (top), or $\beta_2$m bound to unlabeled HC and MART1 (bottom). Additional peaks in the spectra collected at 800 MHz are largely due to protein degradation and do not affect the chemical shifts corresponding to the protein backbone. All data were collected with identical buffer conditions (20 mM sodium phosphate, pH 7.2, and 150 mM NaCl) and at RT (298-300 K).

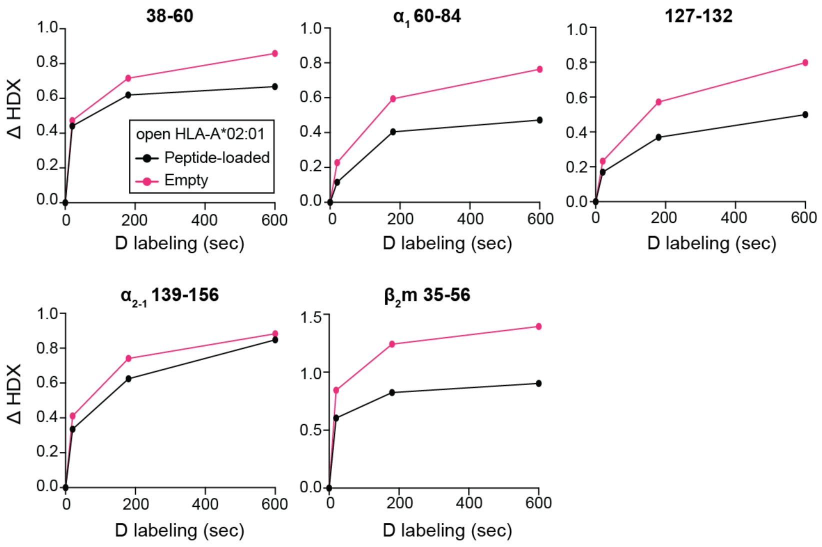


**Figure S4. Percent deuterium uptake resolved to individual peptide fragments.** Peptide segments of 38-60, $\alpha_1$ 60-84, 127-132, $\alpha_{2\text{-}1}$ 139-156, and $\beta_2$m 35-56 are plotted for each exposure time (0, 20, 180, and 600s). The plots reveal the local HDX profiles of HLA-A*02:01 for the states of peptide-loaded (black, refolded with KILGFVFJV) and empty (pink, 40-minute UV irradiation at 4$^{o}$C).

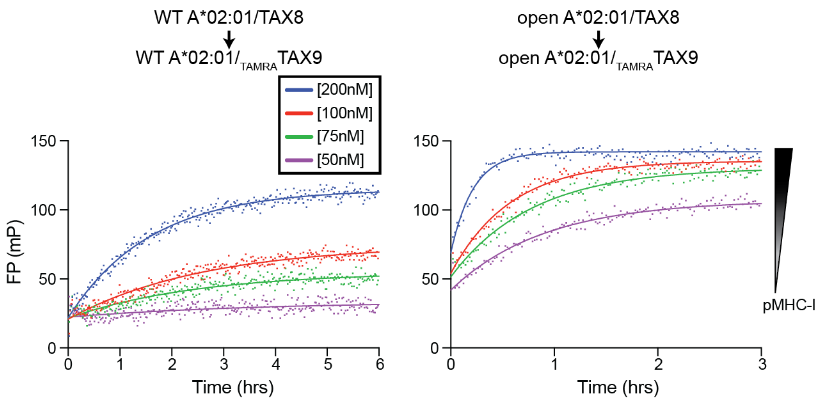


**Figure S5. Binding of $_{TAMRA}$TAX9 by the WT or open HLA-A*02:01/TAX8.** Peptide exchange measured by fluorescence polarization (mP) of 40nM $_{TAMRA}$TAX9 as a function of the WT or open HLA-A*02:01/TAX8 concentrations. Individual traces were fit to a monoexponential association model to determine apparent rate constants $K_{assoc.}$ plotted in **Fig. 4D**. Results of three replicates (mean) are plotted.

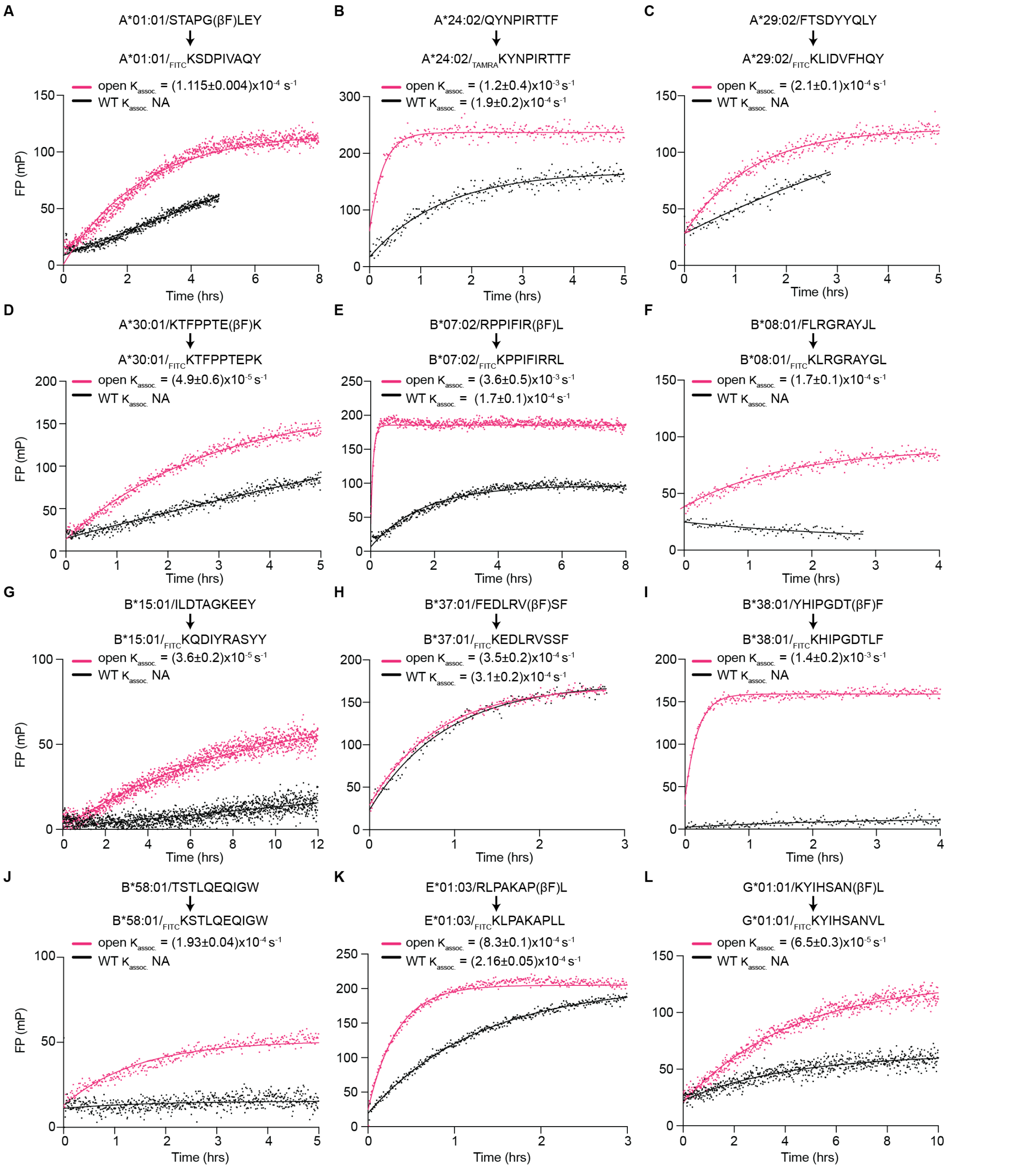


**Figure S6. Peptide exchange kinetics of the open vs. WT HLA allotypes. A-L,** The association profiles of fluorophore-conjugated peptides $_{\text{FITC}}$KSDPIVAQY, $_{\text{TAMRA}}$KYNPIRTTF, $_{\text{FITC}}$KLIDVFHQY, $_{\text{FITC}}$KTFPPTEPK, $_{\text{FITC}}$KPPIFIRRL, $_{\text{FITC}}$KLRGRAYGL, $_{\text{FITC}}$KQDIYRASYY, $_{\text{FITC}}$KEDLRVSSF, $_{\text{FITC}}$KHIPGDTLF, $_{\text{FITC}}$KSTLQEQIGW, $_{\text{FITC}}$KLPAKAPLL, and $_{\text{FITC}}$KYIHSANVL to the open (pink) and WT (black) HLA- **A.** A*01:01/STAPG(βF)LEY, **B.** A*24:02/QYNPIRTTF, **C.** A*29:02/FTSDYYQLY, **D.** A*30:01/KTFPPTE(βF)K, **E.** B*07:02/RPPIFIR(βF)L, **F.** B*08:01/FLRGRAYJL, **G.** B*15:01/ILDTAGKEEY, **H.**

B*37:01/FEDLRV(βF)SF, **I.** B*38:01/YHIPGDT(βF)F, **J.** B*58:01/TSTLQEQIGW, **K.** E*01:03/RLPAKAP(βF)L, and **L.** G*01:01/KYIHSAN(βF)L. The data were fitted to a monoexponential association model to determine apparent rate constants $K_{assoc.}$ NA means the $K_{assoc.}$ cannot be fitted. Results of three replicates (mean ± σ) are plotted**.**

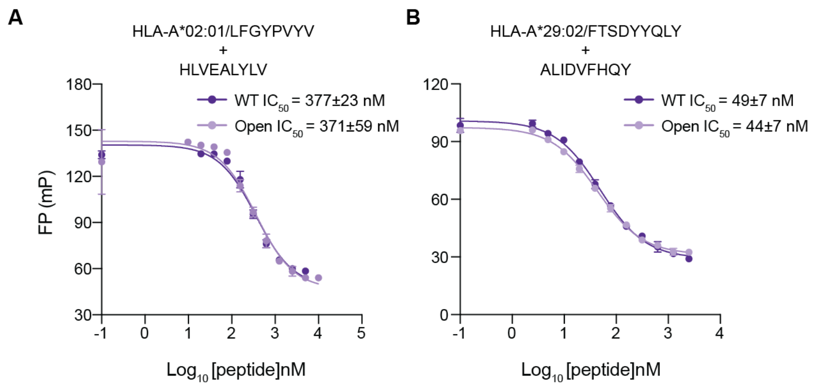


**Figure S7. Selected T1D epitopes demonstrate the same IC50 profiles for the WT or open MHC-I.** The $IC_{50}$ profiles extracted from the association profiles of **A.** $_{TAMRA}$KLFGYPVYV binding to HLA-A*02:01/TAX8 and **B.** $_{FITC}$KLIDVFHQY binding to HLA-A*29:02/FTSDYYQLY in a concentration gradient of a competitor HLVEALYLV and ALIDVFHQY peptides, respectively. $IC_{50}$ values were determined by fitting a log(inhibitor) vs. response (three parameters) curve. Results of three replicates (mean ± $\sigma$) are plotted.

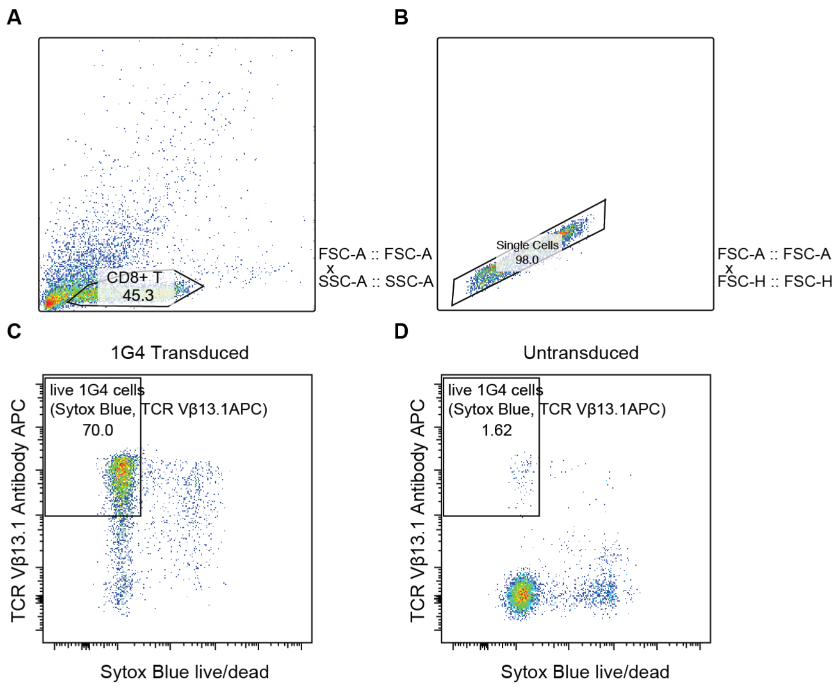


**Figure S8. Flow cytometry gating strategy of CD8+ T cells transduced with 1G4. A-B.** Previously transduced or non-transduced primary human CD8+ T cells were thawed and recovered before tetramer staining. Cells were sorted by side and forward scatter (**A.** SSC-A and FSC-A) followed by single cell isolation (**B.** FSC-A versus FSC-H plot). **C-D.** Gating for live cells was determined by Sytox blue staining, and transduction efficiency was determined by staining with an anti-Vβ13.1-APC antibody (Miltenyi Biotec). Gates are shown in black, and the percentages of events are gated in parentheses. The acquisition was performed on CytoFLEX LX (Beckman Coulter), and the data were analyzed by FlowJo v10.8.1.

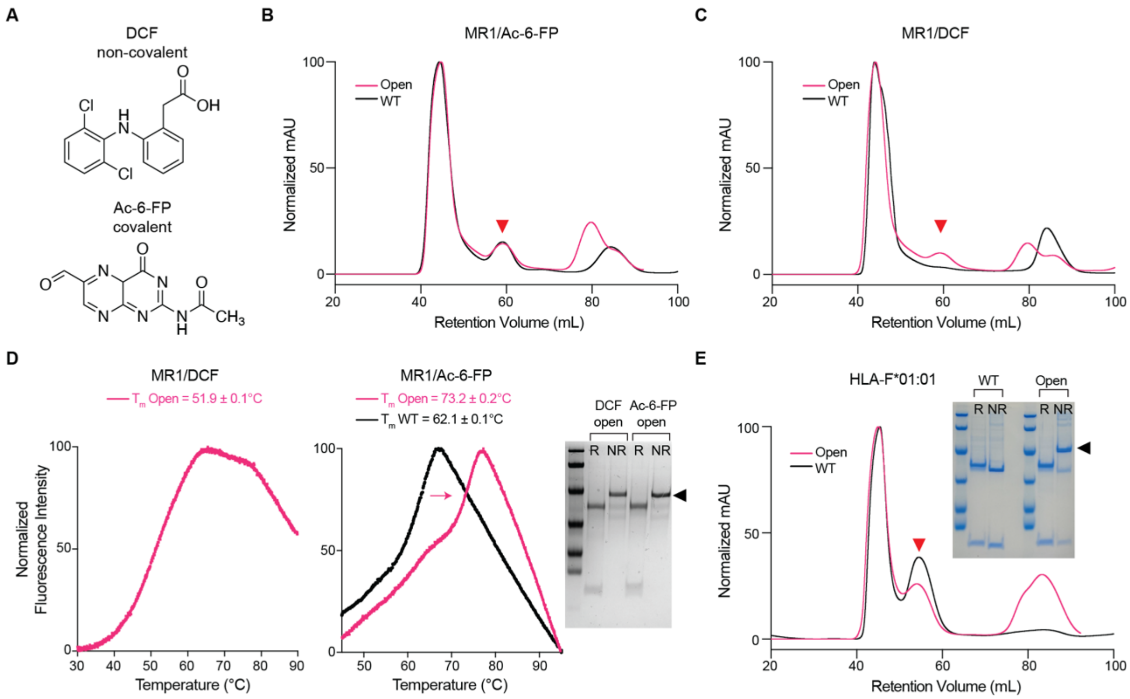


**Figure S9. Disulfide-engineered open MR1 and HLA-F*01:01 molecules form stable protein complexes. A.** Chemical structures of MR1 ligands DCF and Ac-6-FP. **B-C.** SEC traces of the WT (black) and open (pink) MR1 C262S refolded with **B.** Ac-6-FP and **C.** DCF. The triangle arrowhead indicates the protein complexes. **D.** Melting temperature ($T_m$, ºC) obtained from DSF of the WT (black) and open (pink) MR1 C262S loaded with DCF or Ac-6-FP, which are further confirmed by SDS/PAGE analysis in reduced (R) or non-reduced (NR) conditions. Results of three technical replicates (mean ± $\sigma$) are plotted. **E.** SEC traces of the WT (black) and open (pink) HLA-F*01:01/$\beta_2$m. The triangle arrowheads indicate the complex peaks, which are further confirmed by SDS/PAGE analysis in reduced (R) or non-reduced (NR) conditions.

**Supplemental Tables**

| #Peptide | Peptide sequence | Wild type | | Open | |
|---|---|---|---|---|---|
| | | $T_m$ (°C) | σ (±°C) | $T_m$ (°C) | σ (±°C) |
| 1 | KLVVVGACGV | 48.81 | 0.11 | 52.19 | 0.09 |
| 2 | LLGRNSFEVHV | 43.42 | 0.38 | 48.66 | 0.21 |
| 3 | KLVVVGAAGV | 48.06 | 0.11 | 51.65 | 0.12 |
| 4 | RLIRVEGNLRV | 53.37 | 0.25 | 53.33 | 0.17 |
| 5 | KLVVVGASGV | 46.22 | 0.05 | 49.86 | 0.12 |
| 6 | ALNNMFCQLA | 44.42 | 0.11 | 51.07 | 0.40 |
| 7 | ILNREIDFA | 48.69 | 0.07 | 53.00 | 0.18 |
| 8 | KTYPVQLWV | 49.35 | 0.11 | 54.49 | 0.35 |
| 9 | GLAPPQHRI | 44.05 | 0.15 | 49.20 | 0.32 |
| 10 | YMFNSSCMGGM | 48.02 | 0.09 | 53.38 | 0.19 |
| 11 | LLGRNSFEMRV | 46.24 | 0.07 | 50.78 | 0.13 |
| 12 | GLAPPQRLIRV | 50.73 | 0.11 | 53.77 | 0.16 |
| 13 | YLSTDVGFCT | 48.30 | 0.11 | 53.88 | 0.19 |
| 14 | VLMGHVAAVG | 58.50 | 0.77 | 57.73 | 0.05 |
| 15 | YLDSGIHFG | 49.90 | 0.13 | 52.78 | 0.47 |
| 16 | KILNREIDFAI | 46.89 | 0.13 | 52.79 | 0.10 |
| 17 | YMCNSSCMGV | 60.98 | 3.22 | 58.62 | 0.21 |
| 18 | YSSGFCNIAV | 43.73 | 0.06 | 49.31 | 0.16 |
| 19 | LLVRNSFEV | 53.93 | 0.29 | 57.38 | 0.11 |
| 20 | ILWRQDIHLGV | 53.91 | 0.08 | 57.55 | 0.04 |
| 21 | MFCQLAKTYPV | 43.67 | 0.23 | 54.24 | 0.36 |
| 22 | LLVRNSFEVRV | 44.32 | 0.06 | 51.30 | 0.22 |
| 23 | IDILWRQDIHL | 44.83 | 0.06 | 51.71 | 0.61 |
| 24 | KLVVVGADGV | 45.08 | 0.11 | 49.77 | 0.09 |
| 25 | GMNWRPILTI | 42.32 | 0.33 | 50.04 | 0.07 |
| 26 | ILCATYVKV | 57.41 | 0.36 | 59.44 | 0.12 |
| 27 | LLGRNSFEVLV | 42.80 | 0.24 | 48.85 | 0.20 |
| 28 | LLDILDTAGL | 40.96 | 0.05 | 49.75 | 0.09 |
| 29 | CLLDILDTAGL | 46.69 | 0.04 | 54.21 | 0.22 |
| 30 | FSGEYIPTV | 56.01 | 0.51 | 58.30 | 0.05 |
| 31 | RPLAWGNINL | 44.02 | 0.12 | 48.17 | 0.22 |
| 32 | ILNREIDFAI | 44.95 | 0.08 | 52.01 | 0.14 |
| 33 | GLKDLLNPI | 53.70 | 0.24 | 54.58 | 0.29 |

| | | | | | |
|---|---|---|---|---|---|
| 34 | YLDSGIHCGA | 51.99 | 0.38 | 54.34 | 0.20 |
| 35 | FCQLAKTYPV | 50.19 | 0.27 | 52.29 | 0.10 |
| 36 | ILWRQDIHL | 55.74 | 0.37 | 58.83 | 0.02 |
| 37 | GLVDEQQEV | 54.49 | 0.30 | 54.61 | 0.26 |
| 38 | NLLVRNSFEV | 47.57 | 0.22 | 49.85 | 0.20 |
| 39 | KLVVVGAVGV | 49.24 | 0.17 | 52.59 | 0.12 |
| 40 | KILCATYVKV | 47.90 | 0.44 | 51.72 | 0.09 |
| 41 | YLSTDVGFCTL | 50.29 | 0.25 | 54.63 | 0.10 |
| 42 | FMKQMNDAL | 43.99 | 0.50 | 49.62 | 0.04 |
| 43 | YLDSGIHFGA | 54.43 | 0.18 | 56.41 | 0.15 |
| 44 | GLAPPQHLTRV | 55.33 | 0.93 | 57.39 | 0.07 |
| 45 | ALNNMFCQL | 51.43 | 0.78 | 54.62 | 0.51 |
| 46 | LLGRNSFEM | 47.91 | 0.29 | 51.31 | 0.07 |
| 47 | GLKDLLNPIGV | 48.99 | 0.29 | 52.79 | 0.28 |
| 48 | CQLAKTYPV | 55.84 | 1.18 | 62.52 | 0.08 |
| 49 | VLHECNSSYI | 46.60 | 0.20 | 53.12 | 0.19 |
| 50 | KLVVVGAGCV | 47.64 | 0.13 | 49.39 | 0.09 |
| TAX9 | LLFGYPVYV | 65.06 | 0.18 | 63.74 | 0.10 |
| P29 | YPNVNIHNF | 41.99 | 0.15 | 48.72 | 0.07 |
| TAX9 Refolded | LLFGYPVYV | 64.20 | 0.19 | 64.44 | 0.06 |

**Table S1. Thermal stabilities for Cancer Genome Atlas (TCGA) epitope library determined by DSF**. $T_m$ of individual peptides from the TCGA epitope library loaded on WT or open HLA-A*02:01 were measured via peptide exchange in triplicates. The high-affinity HLA-A*02:01-restricted TAX9 peptide and refolded TAX9/A02 molecules were used as positive controls, and the irrelevant peptide p29 was used as a negative control.

| Classical HLA Alleles | |
|---|---|
| HLA-A Supertypes | |
| Allele | Protein Sequence |
| A*02:01 | M**G**SHSMRYFFTSVSRPGRGEPRFIAVGYVDDTQFVRFDSDAASQRMEPRAPWIEQEGPEYWDGETRKVKAHSQTHRVDLGTLRGYYNQSEAGSHTVQRMYGCDVGSDWRFLRGYHQYAYDCKDYIALKEDLRSWTAADMAAQTTKHKWEAAHVAEQLRAYLEGTCVEWLRRYLENGKETLQRTDAPKTHMTHHAVSDHEATLRCWALSFYPAEITLTWQRDGEDQTQDTELVETRPAGDGTFQKWAAVVVPSGQEQRYTCHVQHEGLPKPLTLRWEPGSLHHILDAQKMVWNHR |
| A*01:01 | MAS**G**SHSMRYFFTSVSRPGRGEPRFIAVGYVDDTQFVRFDSDAASQKMEPRAPWIEQEGPEYWDQETRNMKAHSQTDRANLGTLRGYYNQSEDGSHTIQIMYGCDVGPDGRFLRGYRQDAYDCKDYIALNEDLRSWTAADMAAQITKRKWEAVHAAEQRRVYLEGRCVDGLRRYLENGKETLQRTDPPKTHMTHHPISDHEATLRCWALGFYPAEITLTWQRDGEDQTQDTELVETRPAGDGTFQKWAAVVVPSGEEQRYTCHVQHEGLPKPLTLRWELSSQPGSLHHILDAQKMVWNHR |
| A*24:02 | MAS**G**SHSMRYFSTSVSRPGRGEPRFIAVGYVDDTQFVRFDSDAASQRMEPRAPWIEQEGPEYWDEETGKVKAHSQTDRENLRIALRYYNQSEAGSHTLQMMFGCDVGSDGRFLRGYHQYAYDCKDYIALKEDLRSWTAADMAAQITKRKWEAAHVAEQQRAYLEGTCVDGLRRYLENGKETLQRTDPPKTHMTHHPISDHEATLRCWALGFYPAEITLTWQRDGEDQTQDTELVETRPAGDGTFQKWAAVVVPSGEEQRYTCHVQHEGLPKPLTLRWEPSSQPGSLHHILDAQKMVWNHR |
| A*29:02 | M**G**SHSMRYFTTSVSRPGRGEPRFIAVGYVDDTQFVRFDSDAASQRMEPRAPWIEQEGPEYWDLQTRNVKAQSQTDRANLGTLRGYYNQSEAGSHTIQMMYGCDVGSDGRFLRGYRQDAYDCKDYIALNEDLRSWTAADMAAQITQRKWEAARVAEQLRAYLEGTCVEWLRRYLENGKETLQRTDAPKTHMTHHAVSDHEATLRCWALSFYPAEITLTWQRDGEDQTQDTELVETRPAGDGTFQKWASVVVPSGQEQRYTCHVQHEGLPKPLTLRWEPGSLHHILDAQKMVWNHR |
| A*30:01 | M**G**SHSMRYFSTSVSRPGSGEPRFIAVGYVDDTQFVRFDSDAASQRMEPRAPWIEQERPEYWDQETRNVKAQSQTDRVDLGTLRGYYNQSEAGSHTIQIMYGCDVGSDGRFLRGYEQHAYDCKDYIALNEDLRSWTAADMAAQITQRKWEAARWAEQLRAYLEGTCVEWLRRYLENGKETLQRTDPPKTHMTHHPISDHEATLRCWALGFYPAEITLTWQRDGEDQTQDTELVETRPAGDGTFQKWAAVVVPSGEEQRYTCHVQHEGLPKPLTLRWELGSLHHILDAQKMVWNHR |
| HLA-B Supertypes | |
| Allele | Protein Sequence |

| | |
|---|---|
| B*07:02 | MGSHSMRYFYTSVSRPGRGEPRFISVGYVDDTQFVRFDSDAASPREEPRAPWIEQEGPE<br>YWDRNTQIYKAQAQTDRESLRNLRGYYNQSEAGSHTLQSMYGCDVGPDGRLLRGHDQY<br>AYDCKDYIALNEDLRSWTAADTAAQITQRKWEAAREAEQRRAYLEGECVEWLRRYLENG<br>KDKLERADPPKTHVTHHPISDHEATLRCWALGFYPAEITLTWQRDGEDQTQDTELVETRP<br>AGDRTFQKWAAVVVPSGEEQRYTCHVQHEGLPKPLTLRWEPSSQSGSLHHILDAQKMV<br>WNHR |
| B*08:01 | MASGSHSMRYFDTAMSRPGRGEPRFISVGYVDDTQFVRFDSDAASPREEPRAPWIEQE<br>GPEYWDRNTQIFKTNTQTDRESLRNLRGYYNQSEAGSHTLQSMYGCDVGPDGRLLRGH<br>NQYAYDCKDYIALNEDLRSWTAADTAAQITQRKWEAARVAEQDRAYLEGTCVEWLRRYL<br>ENGKDTLERADPPKTHVTHHPISDHEATLRCWALGFYPAEITLTWQRDGEDQTQDTELVE<br>TRPAGDRTFQKWAAVVVPSGEEQRYTCHVQHEGLPKPLTLRWEPSSQSGSLHHILDAQK<br>MVWNHR |
| B*15:01 | MGSHSMRYFYTAMSRPGRGEPRFIAVGYVDDTQFVRFDSDAASPRMAPRAPWIEQEGP<br>EYWDRETQISKTNTQTYRESLRNLRGYYNQSEAGSHTLQRMYGCDVGPDGRLLRGHDQ<br>SAYDCKDYIALNEDLSSWTAADTAAQITQRKWEAAREAEQWRAYLEGLCVEWLRRYLEN<br>GKETLQRADPPKTHVTHHPISDHEATLRCWALGFYPAEITLTWQRDGEDQTQDTELVETR<br>PAGDRTFQKWAAVVVPSGEEQRYTCHVQHEGLPKPLTLRWEPSSQSGSLHHILDAQKM<br>VWNHR |
| B*37:01 | MGSHSMRYFHTSVSRPGRGEPRFISVGYVDDTQFVRFDSDAASPRTEPRAPWIEQEGPE<br>YWDRETQISKTNTQTYREDLRTLLRYYNQSEAGSHTIQRMSGCDVGPDGRLLRGYNQFA<br>YDCKDYIALNEDLSSWTAADTAAQITQRKWEAARVAEQDRAYLEGTCVEWLRRYLENGK<br>ETLQRADPPKTHVTHHPISDHEATLRCWALGFYPAEITLTWQRDGEDQTQDTELVETRPA<br>GDRTFQKWAAVVVPSGEEQRYTCHVQHEGLPKPLTLRWEPGSLHHILDAQKMVWNHR |
| B*38:01 | MGSHSMRYFYTSVSRPGRGEPRFISVGYVDDTQFVRFDSDAASPREEPRAPWIEQEGPE<br>YWDRNTQISKTNTQTYRENLRIALRYYNQSEAGSHTLQRMYGCDVGPDGRLLRGHNQFA<br>YDCKDYIALNEDLSSWTAADTAAQITQRKWEAARVAEQLRTYLEGTCVEWLRRYLENGK<br>ETLQRADPPKTHVTHHPISDHEATLRCWALGFYPAEITLTWQRDGEDQTQDTELVETRPA<br>GDRTFQKWAAVVVPSGEEQRYTCHVQHEGLPKPLTLRWEPGSLHHILDAQKMVWNHR |
| B*58:01 | MGSHSMRYFYTAMSRPGRGEPRFIAVGYVDDTQFVRFDSDAASPRTEPRAPWIEQEGP<br>EYWDGETRNMKASAQTYRENLRIALRYYNQSEAGSHIIQRMYGCDLGPDGRLLRGHDQS<br>AYDCKDYIALNEDLSSWTAADTAAQITQRKWEAARVAEQLRAYLEGLCVEWLRRYLENG<br>KETLQRADPPKTHVTHHPVSDHEATLRCWALGFYPAEITLTWQRDGEDQTQDTELVETR<br>PAGDRTFQKWAAVVVPSGEEQRYTCHVQHEGLPKPLTLRWEPGSLHHILDAQKMVWNH<br>R |
| HLA-Ib & Nonclassical Alleles | |
| Allele | Protein Sequence |

| | |
|---|---|
| E*01:03 | M**G**SHSLKYFHTSVSRPGRGEPRFISVGYVDDTQFVRFDNDAASPRMVPRAPWMEQEGS<br>EYWDRETRSARDTAQIFRVNLRTLRGYYNQSEAGSHTLQWMHGCELGPDGRFLRGYEQ<br>FAYD**C**KDYLTLNEDLRSWTAVDTAAQISEQKSNDASEAEHQRAYLEDTCVEWLHKYLEK<br>GKETLLHLEPPKTHVTHHPISDHEATLRCWALGFYPAEITLTWQQDGEGHTQDTELVETR<br>PAGDGTFQKWAAVVVPSGEEQRYTCHVQHEGLPEPVTLRWEPGSGGGLNDIFEAQKIE<br>WHE |
| G*01:01 | M**G**SHSMRYFSAAVSRPGRGEPRFIAMGYVDDTQFVRFDSDSASPRMEPRAPWVEQEG<br>PEYWEEETRNTKAHAQTDRMNLQTLRGYYNQSEASSHTLQWMIGCDLGSDGRLIRGYE<br>RYAYD**C**KDYLALNEDLRSWTAADTAAQISKRKSEAANVAEQRRAYLEGTCVEWLHRYLE<br>NGKEMLQRADPPKTHVTHHPVFDYEATLRCWALGFYPAEIILTWQRDGEDQTQDVELVE<br>TRPAGDGTFQKWAAVVVPSGEEQRYTCHVQHEGLPEPLMLRWKQGSLHHILDAQKMV<br>WNHR |
| F*01:01 | M**G**SHSLRYFSTAVSRPGRGEPRYIAVEYVDDTQFLRFDSDAAIPRMEPREPWVEQEGPQ<br>YWEWTTGYAKANAQTDRVALRNLLRRYNQSEAGSHTLQGMNGCDMGPDGRLLRGYHQ<br>HAYD**C**KDYISLNEDLRSWTAADTVAQITQRFYEAEEYAEEFRTYLEGECLELLRRYLENGK<br>ETLQRADPPKAHVAHHPISDHEATLRCWALGFYPAEITLTWQRDGEEQTQDTELVETRPA<br>GDGTFQKWAAVVVPSGEEQRYTCHVQHEGLPQPLILRWEQSPQPTIPIGSLHHILDAQKM<br>VWNHR |
| MR1<br>C262S | M**R**THSLRYFRLGVSDPIHGVPEFISVGYVDSHPITTYDSVTRQKEPRAPWMAENLAPDHW<br>ERYTQLLRGWQQMFKVELKRLQRHYNHSGSHTYQRMIGCELLEDGSTTGFLQYAYD**C**Q<br>DFLIFNKDTLSWLAVDNVAHTIKQAWEANQHELLYQKNWLEEECIAWLKRFLEYGKDTLQ<br>RTEPPLVRVNRKETFPGVTALFCKAHGFYPPEIYMTWMKNGEEIVQEIDYGDILPSGDGTY<br>QAWASIELDPQSSNLYSCHVEHSGVHMVLQVPGSLHHILDAQKMVWNHR |
| CD1d | M**A**EVPQRLFPLRSLQISSFANSSWTRTDGLAWLGELQTHSWSNDSDTVRSLKPWSQ<br>GTFSDQQWETLQHIFRVYRSSFTRDVKEFAKMLRLSYPLELQVSAGCEVHPGNASN<br>NFFHVAFQ**C**KDILSFQGTSWEPTQEAPLWVNLAIQVLNQDKWTRETVQWLLNGTCP<br>QFVSGLLESGKSELKKQVKPKAWLSRGPSPGPGRLLLVCHVSGFYPKPVWVKWMR<br>GEQEQQGTQPGDILPNADETWYLRATLDVVAGEAAGLSCRVKHSSLEGQDIVLYWG<br>GGGGLNDIFEAQKIEWHE |
| $\beta_2$m | MIQRTPKIQVYSRHPAENGKSNFLNCYVSGF**C**PSDIEVDLLKNGERIEKVEHSDLSFSKDW<br>SFYLLYYTEFTPTEKDEYACRVNHVTLSQPKIVKWDRDM |

**Table S2. A summary of open MHC-I and $\beta_2$m sequences used in the study.** Below are the protein sequences for the representative alleles from each HLA supertype. Cysteine mutations are colored in red.

| Peptide Name | HLA Allotype | Sequence |
|---|---|---|
| FITC-A01 | A*01:01 | FITC-KSDPIVAQY |
| TAMRA-TAX9 | A*02:01 | TAMRA-KLFGYPVYV |
| TAMRA-PHOX2B | A*24:02 | TAMRA-KYNPIRTTF |
| FITC-A29 | A*29:02 | FITC-KLIDVFHQY |
| FITC-A30 | A*30:01 | FITC-KTFPPTEPK |
| FITC-B07 | B*07:02 | FITC-KPPIFIRRL |
| FITC-B08 | B*08:01 | FITC-KLRGRAYGL |
| FITC-B38 | B*38:01 | FITC-KHIPGDTLF |
| FITC-B37 | B*37:01 | FITC-KEDLRVSSF |
| FITC-B58 | B*58:01 | FITC-KSTLQEQIGW |
| FITC-B15 | B*15:01 | FITC-KQDIYRASYY |
| FITC-E01 | E*01:03 | FITC-KLPAKAPLL |
| FITC-G01 | G*01:01 | FITC-KYIHSANVL |

**Table S3. A summary of the fluorophore-labeled peptides used in the study.**

| HLA Supertype | HLA Allele | Placeholder Ligand | Ligand Sequence | Melting Temperature | | | |
|---|---|---|---|---|---|---|---|
| | | | | wild-type | | open | |
| | | | | $T_m$ (°C) | σ (±°C) | $T_m$ (°C) | σ (±°C) |
| A01 | A*01:01 | β-A01 | STAPG(βF)LEY | 43.75 | 0.02 | 61.2 | 0.5 |
| A0103 | A*30:01 | β-A30 | KTFPPTE(βF)K | 49.18 | 0.08 | 51.05 | 0.09 |
| A0124 | A*29:02 | SARS P44 | FTSDYYQLY | 53.97 | 0.07 | 53.46 | 0.08 |
| A02 | A*02:01 | TAX8 | LFGYPVYV | 41.6 | 0.2 | 48.79 | 0.07 |
| | | TAX9 | LLFGYPVYV | 52.36 | 0.09 | 52.98 | 0.09 |
| A24 | A*24:02 | Phox2B | QYNPIRTTF | 66.0 | 0.1 | 63.23 | 0.05 |
| B07 | B*07:02 | β-B07 | RPPIFIR(βF)L | 44.8 | 0.1 | 48.5 | 0.3 |
| B08 | B*08:01 | PhotoB08 | FLRGRAYJL | 56.5 | 0.2 | 55.93 | 0.07 |
| B27 | B*38:01 | β-B38 | YHIPGDT(βF)F | 49.4 | 0.2 | 50.7 | 0.1 |
| B44 | B*37:01 | β-B37 | FEDLRV(βF)SF | 49.0 | 0.5 | 47.8 | 0.5 |
| B58 | B*58:01 | TW10 | TSTLQEQIGW | 49.3 | 0.2 | 53.1 | 0.4 |
| B62 | B*15:01 | nRASQ61K | ILDTAGKEEY | 46.8 | 0.1 | 52.5 | 0.1 |
| E | E*01:03 | β-E01 | RLPAKAP(βF)L | 49.2 | 0.1 | 51.1 | 0.1 |
| G | G*01:01 | β-G01 | KYIHSAN(βF)L | 60.4 | 0.1 | 51.0 | 0.2 |
| MR1 | MR1 C262S | Diclofenac | - | - | - | 51.9 | 0.1 |
| | | Ac-6-FP | - | 62.1 | 0.1 | 73.2 | 0.2 |

**Table S4. Summary of the designed placeholder peptides and the $T_m$.** Melting temperatures ($T_m$) were determined for the WT and mutant HLA allotype representatives. Each allotype was refolded with a selected placeholder peptide and its $T_m$ was determined by three technical replicates.
